# Cancer Cell Intrinsic Expression of MHCII Regulates the Immune Microenvironment and Response to Anti-PD-1 Therapy in Lung Adenocarcinoma

**DOI:** 10.1101/2020.01.09.900597

**Authors:** Amber M. Johnson, Bonnie L. Bullock, Alexander J. Neuwelt, Joanna M. Poczobutt, Rachael E. Kaspar, Howard Y. Li, Jeff W. Kwak, Katharina Hopp, Mary C. M. Weiser-Evans, Lynn E. Heasley, Erin L. Schenk, Eric T. Clambey, Raphael A. Nemenoff

## Abstract

MHC class II (MHCII) expression is usually restricted to antigen presenting cells, but can be expressed by cancer cells. We examined the effect of <u>cancer cell-intrinsic</u> MHC class II (csMHCII) expression in lung adenocarcinoma on T cell recruitment to tumors and response to anti-PD-1 therapy. The functional significance of altering csMHCII expression was explored using two orthotopic immunocompetent murine models of non-small cell lung cancer: CMT167 (CMT) and Lewis Lung Carcinoma (LLC). We previously showed that CMT167 tumors are eradicated by anti-PD1 therapy, while LLC tumors are resistant. RNA-seq analysis of cancer cells recovered from tumors revealed that csMHCII correlated with response to anti-PD1 therapy, with immunotherapy-sensitive CMT167 cells being csMHCII positive, while resistant LLC cells were csMHCII negative. To test the functional effects of csMHCII, MHCII expression was altered on the cancer cells through loss- and gain-of-function of CIITA, a master regulator of the MHCII pathway. Loss of CIITA in CMT167 decreased csMHCII, and converted tumors from anti-PD-1-sensitive to anti-PD-1-resistant. This was associated with decreased T cell infiltration, lower levels of Th1 cytokines, increased B cell number and decreased macrophage recruitment. Conversely, overexpression of CIITA in LLC cells resulted in csMHCII *in vitro* and *in vivo*. Enforced expression of CIITA increased T cell infiltration and sensitized tumors to anti-PD-1 therapy. csMHCII expression was also examined in a subset of surgically resected human lung adenocarcinomas by multispectral imaging, provided a survival benefit and positively correlated with T cell infiltration. These studies demonstrate a functional role for csMHCII in regulating T cell infiltration and sensitivity to anti-PD-1.

## Introduction

Lung cancer is the leading cause of cancer-related deaths, with an overall 5-year survival rate of less than 18% [1]. Non-small cell lung cancer (NSCLC) represents 85% of all lung cancers [2] and can be divided into adenocarcinomas and squamous cell carcinomas. During the past 10 years, a paradigm shift has changed the view of cancer from an autonomous cellular disease to a complex system of interactions between cancer cells and the tumor microenvironment (TME) [3, 4]. Interactions between cancer cells and T cells can facilitate or hinder tumor progression. In general, increased tumor T cell infiltration correlates with positive clinical outcome [4, 5], indicating that T cells can inhibit tumor progression. However, cancer cells can mobilize multiple mechanisms to evade immune attack, and immunoevasion is one of the hallmarks of cancer progression [6]. Binding of programmed cell death ligand-1 (PD-L1), expressed on cancer cells and other cells of the TME, to programmed cell death protein-1 (PD-1) expressed on T cells results in inhibition of T cell receptor signaling. Prolonged engagement results in a hypofunctional, “exhausted” T cell state that fails to contain tumor progression. Antibodies against either PD-1 or PD-L1 disrupt this interaction, reinvigorating T cell function and can potentially result in tumor elimination [7, 8]. These agents are FDA-approved for multiple malignancies including NSCLC [9, 10]. However, even in patients with high PD-L1 expression, less than half respond in the first line of therapy [11]. While associations with mutational and antigenic burden, an inflamed TME, and levels of PD-1/PD-L1 expression have been described, definitive mechanisms underlying tumor responsiveness or resistance to anti-PD-1/anti-PD-L1 targeted therapies remain highly sought after [12–14].

One well-established mechanism of immunoevasion is the failure of cancer cells to present tumor antigens. For example, loss of MHC class I (MHCI) contributes to immune evasion by decreasing antigen presentation to CD8^+^ cytotoxic T cells, which recognize and directly kill tumor cells [15]. While MHCI is expressed on all cells, MHC class II (MHCII) expression is usually restricted to antigen presenting cells (APCs). Peptide-loaded MHCII molecules are constitutively expressed on APCs and MHCII antigen presentation is essential for CD4^+^ helper T cell-dependent immune responses [16]. MHCII can also be induced on non-APCs in response to an inflammatory environment and inflammatory cytokines such as IFNγ [17]. Among non-APCs, MHCII expression by cancer cells could potentially have an important role in anti-tumor immunity, as it would afford the potential for direct recognition and engagement of cancer cells presenting tumor neoantigens in the context of MHCII to CD4^+^ helper T cells. In this study we designate expression of MHCII on cancer cells as cancer cell-specific MHCII (csMHCII) to distinguish it from expression on other cells in the TME. In colon cancer, high csMHCII expression is associated with increased survival [18, 19]. Similarly, melanomas with high expression of csMHCII respond better to immune checkpoint therapy blockade [20]. In triple negative breast cancer csMHCII expression was also associated with better progression free survival [21, 22]. While these studies describe a positive correlation between csMHCII and survival or response to therapy, whether csMHCII is functionally important in antigen presentation, regulating the TME or promoting the response to immunotherapy remains unclear.

Our laboratory has developed an immunocompetent orthotopic murine model of lung adenocarcinoma in which murine lung cancer cells are directly implanted into the lungs of syngeneic mice [23–26]. This allows the tumors to develop in the appropriate microenvironment. We previously compared the response of two KRAS mutant lung cancer cell lines, CMT167 and Lewis Lung Carcinoma (LLC), to immune checkpoint inhibitors [24]. CMT167 tumors were strongly inhibited by anti-PD-1 or anti-PD-L1 antibodies, whereas LLC tumors were resistant. The mechanisms underlying this differential response are not well understood. In this study, we present evidence that csMHCII expression positively correlates with response to immunotherapy and test the functional contribution of csMHCII on anti-tumor immunity under baseline and immunotherapy conditions. We further demonstrate that csMHCII expression varies between human lung adenocarcinomas and is positively correlated with T cell infiltration and a survival benefit. These studies emphasize the unique contribution of csMHCII on shaping the TME and influencing the response to immunotherapy.

## Materials and Methods

### Cell Lines and Culture Conditions

CMT167 cells were stably transfected with firefly luciferase as previously described [24]. Luciferase-expressing Lewis Lung Carcinoma (LLC) cells were purchased from Caliper Life Sciences (LL/2-luc-M38). The CMT167 cells harbor a *Kras*^G12V^ mutation, and the LLC harbor a *Kras*^G12C^ mutation [24]. Both cell lines were maintained in DMEM with 4.5 g/L glucose (#10-017-CV, Corning) containing 10% FBS, 100 U/mL penicillin, 100 μg/mL streptomycin, and 500 μg/mL G418 at 37°C in a humidified 5% CO2 atmosphere. Cells were periodically tested for mycoplasma infection, maintained as frozen stocks and cultured for only 2-4 weeks before use in experiments. Authentication of cell lines based on morphology, growth curve analysis, and metastatic phenotype was performed regularly. Human NSCLC cell lines were maintained in RPMI 1640 (#10-040-CV, Corning) containing 10% FBS at 37°C in a humidified 5% CO2 atmosphere. All were KRAS mutant.

### Orthotopic Murine Model and anti-PD-1 Treatment

Wild-type C57BL/6J and green-fluorescent protein (GFP) expressing mice [C57BL/6-132Tg (UBC-GFP) 30Scha/J] were obtained from Jackson Laboratory (Bar Harbor, ME). Mice were bred and maintained in the Center for Comparative Medicine at the University of Colorado Denver. Experiments were performed in 7-12 week-old male mice. All procedures were performed under protocols approved by the Institutional Animal Care and Use Committee at the University of Colorado Denver. Cancer cells were directly implanted into the left lobe of the lung as previously described [23, 24]. For experiments with checkpoint inhibitors, tumor-bearing mice were intraperitoneally injected (8-10mg/kg) with either an IgG2a isotype control antibody (BioXCell #BE0089), or an anti-PD-1 antibody (BioXCell #BE0146) twice weekly starting 7-10 days after injection of cancer cells.

### Transcriptome profiling by RNA sequencing (RNA-seq) of lung cancer cells from tumor-bearing GFP-expression transgenic mice

Cancer cells were injected into GFP expressing transgenic mice as described above. Four weeks after CMT167 and two weeks after LLC tumor implantation, mice were sacrificed and single cell suspensions were prepared as described below. Lung cancer cells were recovered by sorting for GFP-(negative) cells and RNA-seq was performed as previously described [23]. Cell sorting was performed at the University of Colorado Cancer Center Flow Cytometry Shared Resource using an XDP-100 cell sorter (Beckman Coulter).

### Generation of CIITA knockdown cells and CIITA overexpression cells

CMT67 cells were stably transfected using lentivirus with shRNA constructs purchased from the University of Colorado Functional Genomics core: non-targeting control (SHC001V), shCIITA48 (TRCN0000086448), shCIITA49 (TRCN0000086449), shCIITA50 (TRCN0000086450). Selection of stable transductants was carried out with puromycin (2 μg/ml, Sigma). Knockdown efficiency was confirmed by qRT-PCR. LLC-luc cells were stably transfected with CIITA overexpression vector pcDNA3-myc-tagged CIITA (addgene Plasmid #14650) or control pcDNA3.1 (Thermofisher V79020) using Lipofectamine 2000 Reagent (Invitrogen 11668-027). Cells were selected with G418 (500ng/mL, Corning) and individual clones screened for expression by flow cytometry.

### Immunofluorescence

Tissues were fixed in 4% paraformaldehyde. Paraffin embedded samples were cut into 5 μm sections. Sections were dehydrated, immersed in 0.1% Sudan Black B (Sigma 199664-25G) in 70% ethanol for 20 minutes, washed in TBST, incubated in citrate antigen retrieval solution at 100°C for 2 hours, washed in 0.1M glycine/TBST (Sigma G-8898) for 10 minutes, and placed in 10mg/ml sodium Borohydride in Hank’s buffer (Gibco, 14175-095). After blocking with 10% goat serum in equal parts of 5% BSA in TBST and Superblock (ScyTek Laboratories, AAA999) at 4°C overnight, sections were incubated with primary anti-CD3 (Thermo Scientific, MA5-14524) anti-CD4 (eBiosciences #14-9766-82), or anti-CD8 (eBiosciences, #14-0808-82) antibody in equal part solution of 5% BSA in TBST and Superblock for 1 hour at room temperature, washed in TBST, and incubated with secondary goat anti-rabbit IgG Alexa Flour 488 (Invitrogen# A11034) or goat anti rabbit IgG Alexa Fluor 594 (Invitrogen# A11037). Slides were mounted with Vectashield mounting medium with DAPI (Vector Laboratories #H-1200). Positive staining was determined by count per high power field (40X), each tumor count was an average of 5 different views of the tissue, of 3 or more separate tumors, and averaged between two different observers.

### Preparation of single cell suspension and flow cytometry

Mice were sacrificed at 4 weeks (CMT167 tumors) or 3 weeks (LLC tumors) and left lungs containing tumors were excised. Single cell suspensions were prepared as previously described [23–25]. Cells were stained with three panels of antibodies (see **Supplemental Table 1**). For intracellular markers, cells were treated with the Brefeldin A (Biolegend #420601) and Monensin (Biolegend #420701) and a cell stimulation cocktail (PMA/Ionomycin), for 5 hours at 37° C. Cells were then permeabilized using the FOXP3/Transcription Factor Staining Buffer Set (eBioscience #00-5523-00) and stained with intracellular antibodies. Cells were analyzed at the University of Colorado Cancer Center Flow Cytometry Core using a Gallios flow cytometer (Beckman Coulter). All flow cytometry experiments included single stain cell controls, isotype controls for intracellular stains and subjected to post-acquisition compensation using VersaComp Antibody Capture Bead Kit (Beckman Coulter #B22804). Data was analyzed using Kaluza Software (Beckman Coulter). Each measurement represents three separate isolations from lungs of three pooled mice. Gating strategy in **Supplemental Figure S2C**.

### Co-culture of CD4^+^ T lymphocytes and cancer cells, and analysis by flow cytometry

CD4^+^ T cells were isolated using the EasySep Mouse CD4^+^ T-cell isolation kit (Stemcell Technologies). The CD4^+^ T cells were then added in a 1:1 ratio to cultured CMT167 cells, either non-targeting controls (NT-ctrl) or cells silenced for CIITA (CMT shCIITA) that were pretreated with IFNγ (100ng/mL) (Preprotech #315-05) for 48 hours. The cells were centrifuged at 100xg for 5 min. and incubated overnight at 37°C.

### Quantitative Real-time-PCR

Cells were harvested and homogenized in 350 μL of RLT buffer (Qiagen). Total RNA was isolated using an RNAeasy Kit (Qiagen) and reverse transcription was performed on 1 μg of total RNA using qScript cDNA SuperMix (Quanta BioSciences 95048-025). Real-time PCR analysis was conducted in triplicate in a C1000 Touch Thermal Cycler (Bio-Rad) using Power SYBR Green PCR Master Mix (Applied Biosystems 4367659). The relative message levels of each target gene were normalized to β-actin. *β-actin* Fwd: GGCTGTATTCCCCTCCATCG, Rev: CCAGTTGGTAACAATGCCATG. Murine *Ciita* Fwd: TGCGTGTGATGGATGTCCAG, Rev: CCAAAGGGGATAGTGGGTGTC. *H2-Aa* Fwd: ATCGTGGTGGGCACCATCTTCA, Rev: AAGAGGGACACACGCCTTCCTT. *H2-Dma* Fwd: CTCGAAGCATCTACACCAGTG, Rev: TCCGAGAGCCCTATGTTGGG. *Cd74* Fwd: CGCGACCTCATCTCTAACCAT, Rev: ACAGGTTTGGCAGATTTCGGA. *H2-k1* Fwd: GCTGGTGAAGCAGAGAGACTCAG, Rev: GGTGACTTTATCTTCAGGTCTGCT. *H2-d1* Fwd: CTCCGTCCACTGACTCTTAC, Rev: GAGAACTGAGGGCTCTGGATG.

### Western Blot Analysis

Proteins harvested in lysis buffer (50 mM β-glycerophosphate, pH 7.2, 0.5% Triton X-100, 5 mM EGTA, 100 μM sodium orthovanadate, 1mM dithiothreitol, 2 mM MgCl_2_,) with protease inhibitor cocktail (Sigma). Proteins were fractionated by sodium dodecyl sulfate– polyacrylamide gel electrophoresis, electrophoretically transferred to PVDF membranes and blocked for 1 hour in 5% BSA and TBST (Sigma G-8898). Antibodies: Anti-Mouse pSTAT1 (Y701) (Cell Signaling #9167s), Anti-Mouse STAT1 (Cell Signaling #9172S), Anti-Mouse CIITA (abcam #ab49132), β-Actin (Sigma #A5441).

### Spectral flow antibody staining

Single cells suspensions were stained with a panel of antibodies (see **Supplemental Table 1**). Zombie NIR Viability dye was diluted based upon Biolegend’s & Cytek’s suggested protocol before staining samples at room temperature for 15 minutes (Biolegend #423105) in 1.5mL Eppendorf tubes. After incubation, samples were resuspended and subjected by Fc receptor blockade (eBioscience CD16/CD32 #14-0161-86) followed by surface antibody staining with fluorescently labeled-antibodies. All samples were covered in foil and incubated for a total of 30min at room temperature in the dark. Cells were then permeabilized using the FOXP3/Transcription Factor Staining Buffer Set (eBioscience #00-5523-00) at 4°C for 18-24hrs and then subjected to a 2hr incubation at 4°C for intracellular staining. All samples were washed with FAE buffer and filtered through 35μm cell strainer tubes (Falcon 5ml 12×75mm Polystyrene Round-Bottom Tube #352235) before instrument analysis. Experiments included single stain compensation beads for unmixing purposes (Invitrogen eBioscience Ultra Comp Beads #01-2222-42) as well as single stain cell controls for validation and gating strategies. Samples were analyzed at the University of Colorado Cancer Center Flow Cytometry Core on the Cytek Aurora spectral flow cytometer using SpectroFlo, with data visualized using Kaluza Software (Beckman Coulter).

### Patient Studies

Tumor tissues were collected through a protocol approved by the Mayo Clinic Institutional Review Board and obtained from the Mayo Clinic Lung Cancer Repository. Written informed consent was obtained from all patients in accordance with the Declaration of Helsinki. Patients were identified from the tissue repository that underwent curative surgical resection of lung adenocarcinoma between 2004 and 2007 and had available residual tumor specimens. The electronic medical record was reviewed and pertinent clinical data including age at time of surgery, gender, smoking status, pack years, and last follow up date or date of death was extracted. Patients were staged based on surgical findings using the 7th edition of the AJCC TNM system for NSCLC. For cancer specific survival, patients were censored at the date of last follow up or death from causes other than lung cancer. Formalin-fixed paraffin-embedded tissue blocks were sectioned into 5μm slides by the Pathology Research Core (Mayo Clinic, Rochester, MN).

### Tissue multiplex immunohistochemistry

Sections from FFPE blocks were stained using Opal multiplex according to manufacturer’s protocol (PerkinElmer) for DAPI, HLA-DR, CK, CD3 and CD8 at the Human Immune Monitoring Shared Resource at CU Anschutz Medical Campus. Slide scanning was performed on the Vectra 3.0 instrument. 3-5 multispectral regions of interests were selected and analyzed using the inForm Software 2.4. Images were spectrally unmixed, evaluated for staining intensities and morphology, tissue segmented based on tissue markers, cell segmented based on nuclear and membrane markers and phenotypically scored.

### RNA-seq data acquisition

RNA-seq data for 15 human KRAS mutated lung adenocarcinoma was obtained from the Cancer Cell Line Encyclopedia (CCLE) [27].

### Statistical Analysis

Statistical Analyses were performed using the GraphPad Prism 7 or R software. Data are presented as means ± SEM. A one- or two-way ANOVA was used to compare differences in more than two groups. A student’s t-test was used to compare differences between two groups in data with a normal distribution. A Pearson test was used to test correlation. A Log-rank Test were used to compare survival. In all circumstances, p-values ≤0.05 were considered significant.

## Results

### NSCLC Cells Differentially Upregulate an Antigen Presentation Pathway Signature

We hypothesized that the differential response to anti-PD-1 therapy of CMT167 tumors (sensitive) vs. LLC tumors (resistant) that we previously described [24] was an inherent property of the cancer cells, and how they respond to signals from the TME. To define these differences we analyzed changes in gene expression, comparing cancer cells recovered from *in vivo* tumors by fluorescence-activated cell sorting (FACS) with identical cells grown *in vitro*. CMT167 and LLC cells were injected into separate groups of C57BL/6 GFP+ transgenic mice, and the GFP-(negative) population which represents purified cancer cells, was subjected to RNA sequencing [28]. KEGG pathway analysis determined CMT167 cells harvested from primary tumors showed an enhanced antigen presentation signature relative to LLC cells harvested from primary tumors. Among these genes, CMT167 cells recovered from tumors selectively induce MHCII genes and cofactors necessary for processing and presentation, such as CIITA, the master transcriptional regulator of the MHCII pathway, whereas the LLC cells did not (**Figure 1A**). To validate the induction of csMHCII at the protein level, separate single cell suspensions of tumor-bearing lungs grown from C57BL/6 GFP+ mice were analyzed by flow cytometry. Flow cytometric analysis demonstrated that the GFP-(negative) CMT167 cancer cells showed a significant increase of surface MHCII compared to cells cultured *in vitro*. Importantly, csMHCII expression was not observed in LLC tumors (**Figure 1B**). For both tumors, we detected MHCII expression on the GFP+ population, which reflects expression on APCs of the TME (**Supplementary Fig S1A and S1B**).

**Figure 1:**
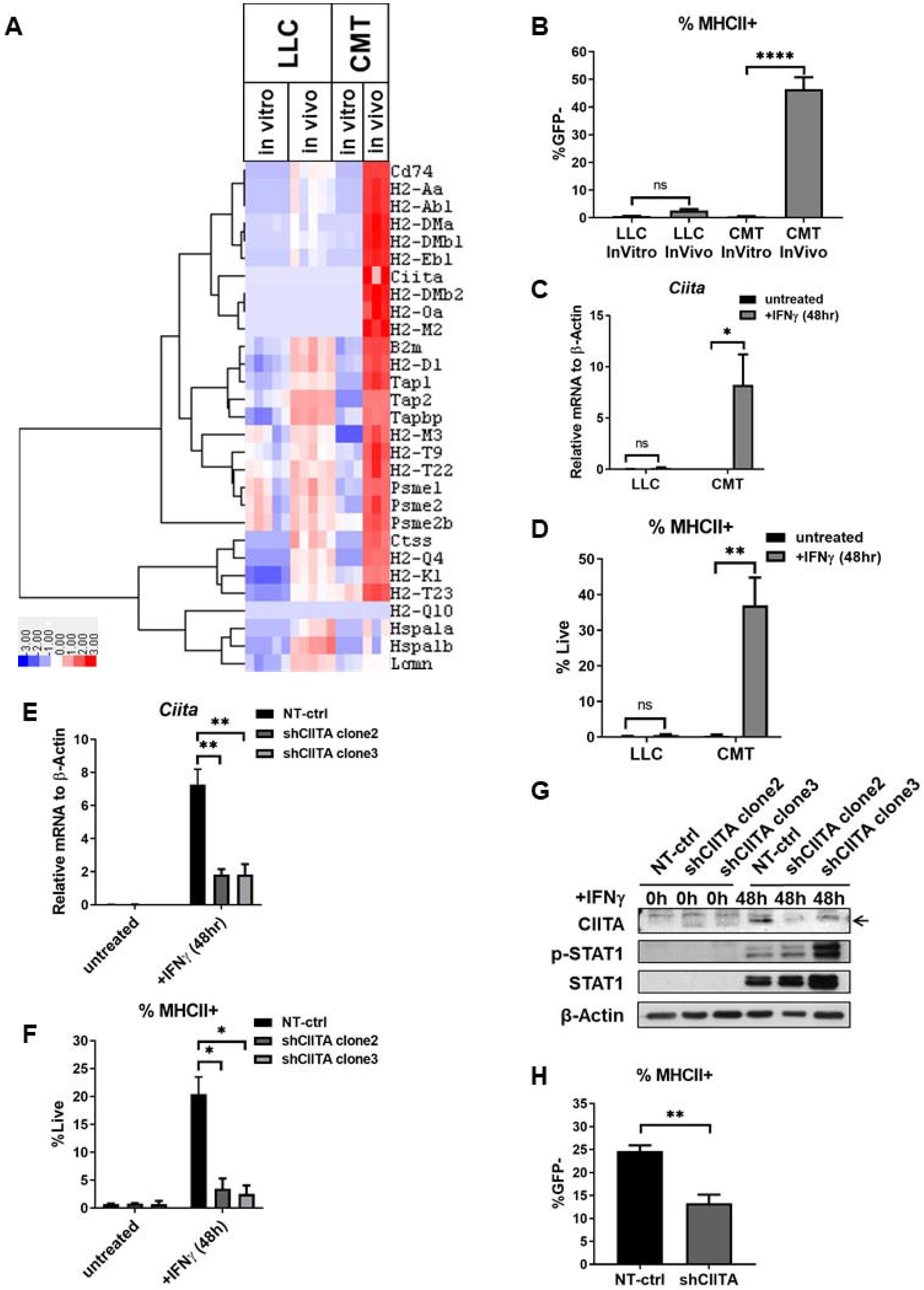
NSCLC Cells Differentially Express an Antigen Presentation Signature. **A)** Heatmap of RNA-seq of MHC associated genes from the antigen processing presentation signature using Kegg Pathway Analysis *in vitro* and *in vivo* of LLC or CMT167. **B)** *Ciita* mRNA expression in LLC and CMT167 cells measured by qRT-PCR (n=3, *p*<.0001****, Unpaired T-test). **C)** %MHCII+ expression measured in LLC and CMT167 cells measured by flow cytometry gated on singlets of live cells (n=3, *p*<.05*, Unpaired T-test). **D)** The %MHCII+ staining measured by flow cytometry gated on singlets of live cells of CMT167 or LLC cells cultured *in vitro* and or CMT167 or LLC tumor cells from C57BL/6 GFP+ mice gated on singlets of live cells of GFP-*in vivo* (n=5, *p*<.01**, One-way ANOVA). **E)** CIITA mRNA expression in CMT167 cells expressing a non-targeting control shRNA (NT-ctrl) or a shRNA for CIITA (shCIITA) untreated or treated with IFNγ for 48hrs measured by qRT-PCR (n=3, *p*<.01**, One-way ANOVA). **F)** The % of MHCII+ staining from live, singlet cells of CMT167 cells expressing a NT-ctrl shRNA or a shRNA for CIITA untreated or treated with IFNγ for 48 hrs measured by flow cytometry (n=3, *p*<.05*, Unpaired T-test). **G)** Western Blot of CIITA, p-STAT1, and total STAT1 in cells cultured in the presence of absence of IFNγ. **H)** Flow cytometric analysis quantifies the % of MCHII+ events focused on live, singlet cells that were GFP-(negative) (n=3, *p*<.01**, Unpaired T-test).

Despite their divergent MHCII induction *in vivo*, both CMT167 and LLC cells expressed low levels of csMHCII expression basally *in vitro*. Since MHCII expression can be induced via IFNγ signaling, we examined whether IFNγ can induce MHCII genes and surface MHCII expression *in vitro*. CMT167 cells treated with IFNγ for 48 hours had increased levels of the MHCII master regulator CIITA, as assessed both by mRNA and protein, as well as increased mRNA expression of other MHCII genes, including *H2-Aa*, *H2-Dma*, and *Cd74* (**Figure 1C, Supplemental Figure S1C and S1D**). This was confirmed by increased level of surface MHCII by flow cytometry (**Figure 1D**). Although LLC cells express IFNγ receptors and upregulate some genes following IFNγ treatment, LLC cells failed to induce MHCII upon IFNγ treatment (**Supplemental S1D**).

### Loss of CIITA decreases MHCII induction

To define the functional role of csMHCII expression, CMT167 cells were transduced with lentiviral constructs encoding three different short hairpins RNA targeting CIITA (shCIITA), the MHCII transactivator, or a non-targeting control short hairpin RNA (NT-ctrl). Since CIITA is a known transcriptional activator of multiple MHCII genes, silencing should disable the whole MHCII antigen presentation pathway while having minimal impact on other pathways [29, 30]. In NT-ctrl CMT167 cells, IFNγ induced *Ciita* mRNA similar to the parental cell lines. However, this induction was impaired in CMT167 transduced with three distinct CIITA shRNAs (**Supplemental Figure S1E**). Similar decreases in cells subjected to knockdown for *Ciita* were also detected in other MHCII pathway genes including *H2-Aa*, *H2-Dma*, and *Cd74* (**Supplemental Figure S1E**). We isolated single-cell clones from the pools of one of the knockdowns, shCIITA(48) to achieve a better CIITA knockdown. Clone2 of shCIITA(48) had decreased CIITA expression, other MHCII genes and surface expression of MHCII protein (**Figure 1E, 1F, and Supplemental Figure S1F**), but showed no alterations in STAT1 signaling or changes induction of MHCI genes in response to IFNγ (**Figure 1G, Supplemental Figure S1G**). In addition, *in vitro* knockdown of CIITA did not alter *Cd274* (*PD-L1)* mRNA induction in response to IFNγ (**Supplemental Figure S1H**). All subsequent *in vivo* experiments were therefore performed with this clone (designated CMT167-shCIITA). To validate knockdown of MHCII on the cancer cells *in vivo*, CMT167-NT-ctrl or CMT167-shCIITA cells were injected into C57BL/6 GFP+ transgenic mice. The CIITA knockdown tumor cells (gated on GFP-(negative) events) showed a decrease in csMHCII expression *in vivo* (**Figure 1H, Supplemental Figure S1I**). The CIITA knockdown did not affect MHCII expression in the TME GFP+ population or MHCI expression on the cancer cells(**Supplemental Figure S1J and S1K**). Thus, CIITA knockdown in CMT167 cells reduces expression of csMHCII with minimal impact on MHCII expression in non-cancer cells of the TME.

### Loss of MHCII decreases CD4^+^ and CD8^+^ tumor infiltration and activation

To examine how csMHCII expression influenced T cell recruitment, infiltrating T cells in CMT167-NT-ctrl tumors compared to CMT167-shCIITA tumors were quantified by immunostaining. CMT167-shCIITA tumors, with reduced csMHCII, had a lower CD3 T cell density (NT-ctrl 139.8±17.96 vs. shCIITA 53.50±8.82), CD4^+^ T cell density (NT-ctrl 83.66±9.903 vs. shCIITA 38.43±7.050) and CD8^+^ T cell density (NT-ctrl 54.53±9.968 vs. shCIITA 27.15±2.605) (**Figure 2A**). Representative images are shown in **Figure 2B**. To determine potential mechanisms to account for the decrease in T cells we isolated RNA from CMT167-NT-ctrl or CMT167-shCIITA whole tumors and performed qRT-PCR for T cell chemoattractants. The shCIITA tumors trended towards decreased levels of *Ifnγ* and *Cxcl9* (**Figure 2C, 2D)**. There were no changes in *Cxcl10*, *Cxcl11*, and a slight decrease in *Cxcl12* (**Figure 2E-2G)**. To determine which cells were producing C*xcl9*, we injected CMT167-NT-ctrl or CMT167-shCIITA into GFP+ mice and then harvested the tumors and recovered the GFP+ by FACS; due to the small size of the tumors we were unable to recover sufficient RNA from the GFP-(negative) population. By qRT-PCR, there was a significant decrease in *Cxcl9* production in GFP+ cells, with no change in *Cxcl10* (**Supplemental Figure S2A, S2B**).

**Figure 2:**
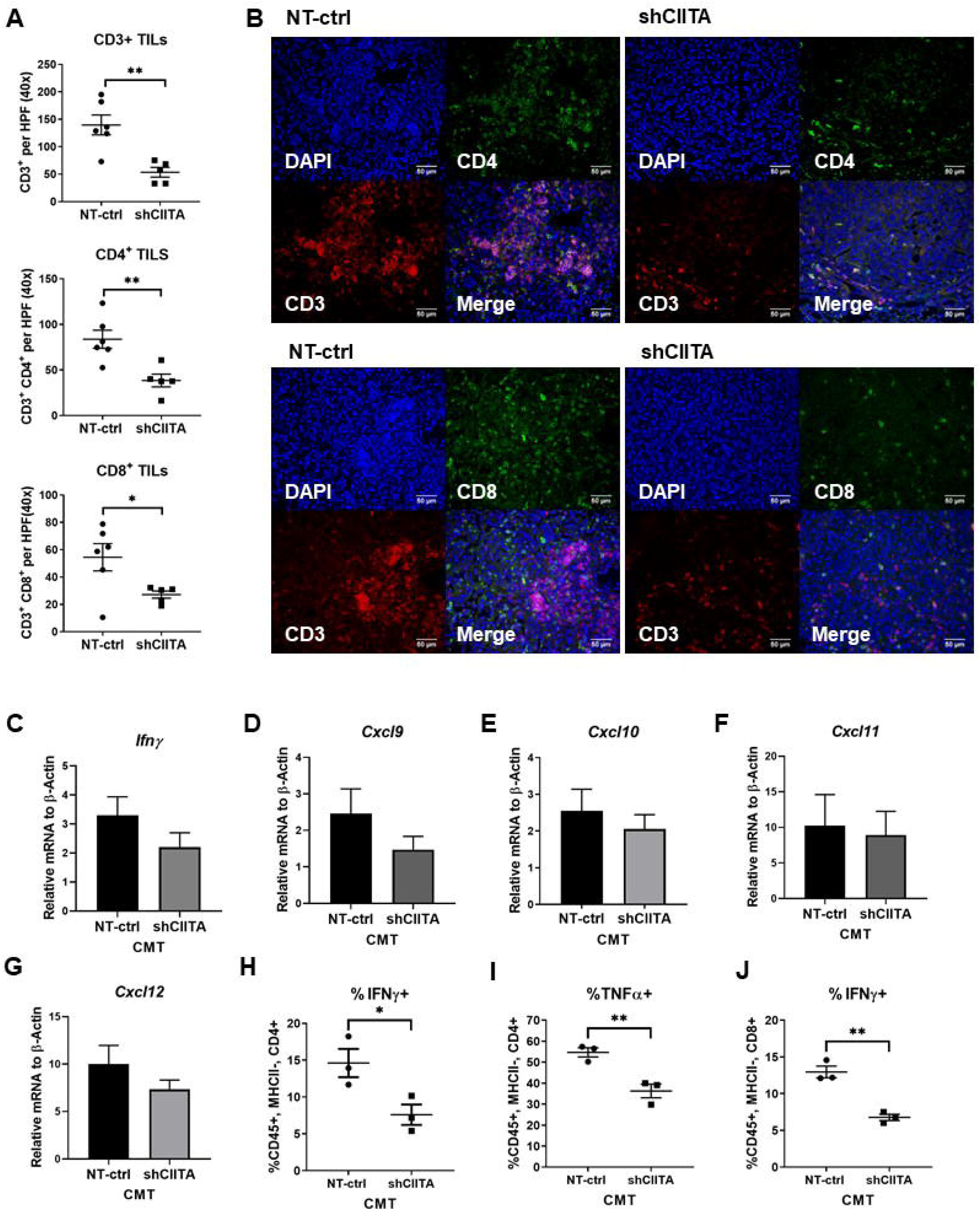
Loss of MHCII decreases CD4^+^ and CD8^+^ tumor infiltration and decreases T cell effector function. **A)** Average Count of immunofluorescence of CD3^+^, CD4^+^,and CD8^+^ positive T cells per high power field (40x) in CMT167-NT-ctrl tumors and CMT167-shCIITA tumors (n=6, n=5, *p*<.05*, *p*<.01**, Unpaired T-test). **B)** Representative pictures of CMT167-NT-ctrl and CMT167-shCIITA tumors IF, DAPI (blue), CD3 (red), and CD4 or CD8 (green). qRT-PCR analysis of CMT167-NT-ctrl and CMT167-shCIITA whole tumors of **C)** Ifnγ **D)** Cxcl9 **E)** Cxcl10 **F)** Cxcl11 and **G)** Cxcl12 (n=4, Unpaired T-test). Flow cytometry on tumor bearing lungs events gated on singlets that are live, CD45+, MHCII-that were either CD4^+^ or CD8^+^; **C)** IFNγ^+^ CD4^+^ T cells, **D)** TNFα^+^ CD4^+^ T cells (n=3, *p*<.05*, *p*<.01**, Unpaired T-test). Percent positive of **H)** IFNγ^+^CD8^+^ T cells in CMT167-NT-ctrl and CMT167-shCIITA tumors (n=3, *p*<.01**, Unpaired T-test).

To understand the functional consequence of csMHCII on effector CD4^+^ T cell function, we examined different CD4^+^ T cell populations and function by flow cytometry. We observed a statistically significant decrease in CD4^+^ Th1 effectors, defined by their expression of IFNγ and TNFα, in CMT167-shCIITA tumors compared to CMT167-NT-ctrl (**Figure 2H, 2I**). csMHCII had no discernable impact on either IL4+ Th2 (**Supplemental Figure S2D**) or IL17+ Th17 cells (**Supplemental Figure S2E**). Tregs, defined by FoxP3 expression, showed a trend toward being increased, but this did not reach statistical significance (**Supplemental Figure S2F**). There were also no changes observed in CD69^+^ and CD44^+^ activation markers among CD4^+^ T cells from tumor bearing lungs (**Supplemental Figure S2G, S2H**). CD8^+^ T cells also had a decrease in effector function as defined by a decrease in IFNγ production, but no significant change in TNFα production (**Figure 2H, Supplemental Figure S2I**) or expression of CD69^+^ and CD44^+^ activation markers (**Supplemental Figure S2J, S2K**). There was no significant change in CTLA4 expression on CD4^+^ or CD8^+^ T cells or PD-1 expression on CD8^+^ T cells. However CD4^+^ T cells had an increase in PD-1 in the CIITA knockdowns (**Supplemental S2L-S2O**). These data demonstrate that loss of csMHCII is associated with reduced anti-tumor Th1 responses and suggests that csMHCII influences the effector functions of the antitumor CD4^+^ T cell response.

### NSCLC cells expressing MHCII directly induce CD4^+^ T cell cytokine production of a Th1 phenotype *in vitro*

To determine if csMHCII expression is functional and can directly activate CD4^+^ T cells, CD4^+^ T cells isolated from lungs of CMT167 tumor-bearing mice were co-cultured overnight with CMT167-NT-ctrl or CMT167-shCIITA cells that had been pretreated with IFNγ for 48 hours to induce MHCII expression. CD4^+^ T cell activation was analyzed by flow cytometry. CD4^+^ T cells co-cultured with CMT167-NT-ctrl cells showed a significant increase in IFNγ and TNFα production, characteristic of a Th1 phenotype, that was not observed in either CD4^+^ T cells cultured alone or with CD4^+^ T cells co-cultured with CMT167-shCIITA cells (**Figure 3A, 3B, Supplemental Figure S2P**). CD4^+^ T cells co-cultured with the CMT167-NT-ctrl cells did not show changes in IL10 production associated with a Th2 phenotype or expression of FoxP3 (**Figure 3C, 3D**). In addition, csMHCII did not influence the frequency of CD69^+^ events (**Figure 3E**). These data provide direct evidence that csMHCII is functional, allowing direct interaction with antitumor CD4^+^ T cells to induce Th1 effector function using an *in vitro* co-culture system.

**Figure 3:**
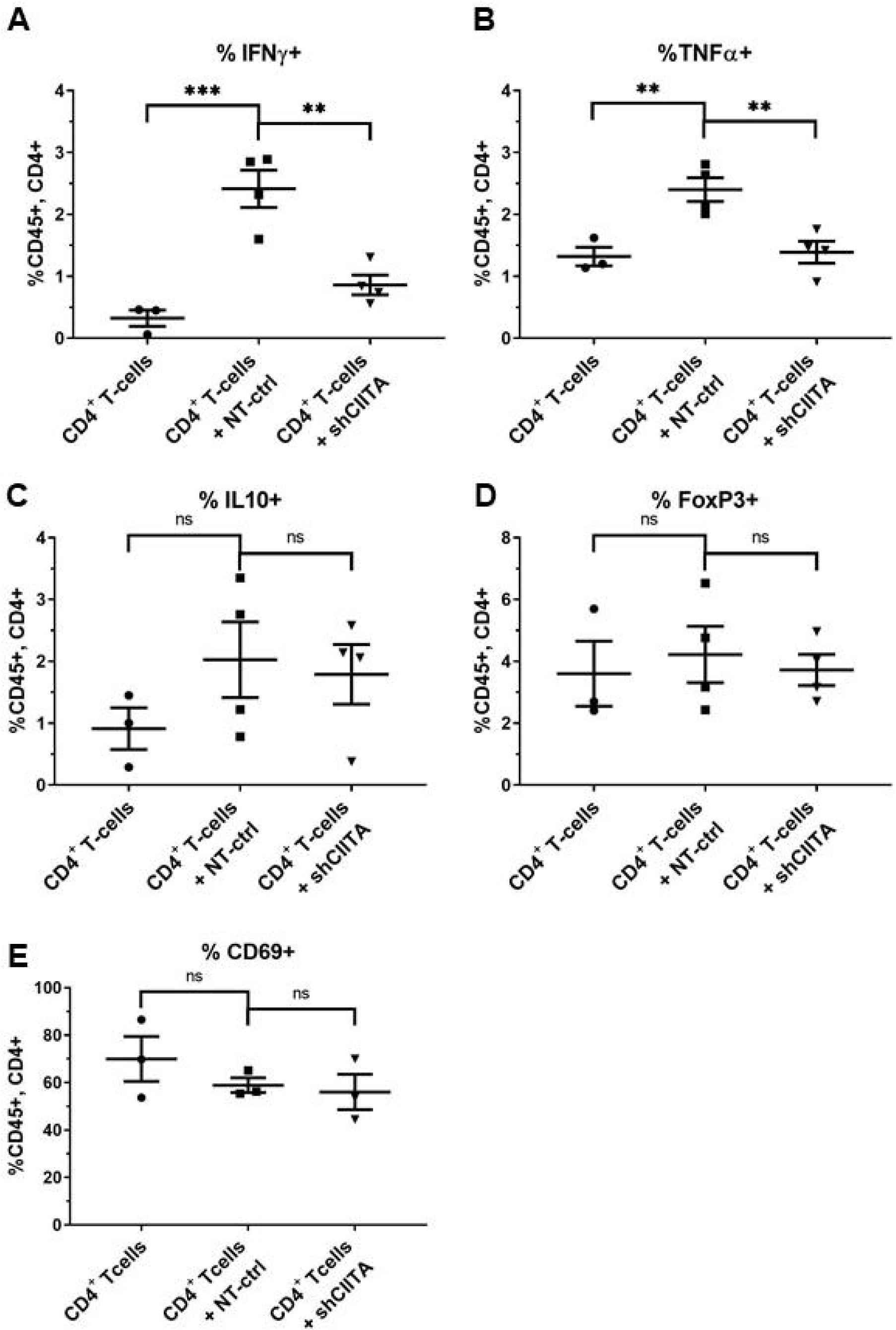
NSCLC cells expressing MHCII directly activate CD4^+^ T cells *in vitro*. Flow cytometric analysis of CD4^+^ T cells isolated from a tumor bearing lung and co-cultured for 24 hours with CMT167-NT-ctrl or CMT167-shCIITA, defined as singlets that are live, CD45+, CD4+. **A)** %IFNγ^+^ CD4^+^ T cells **B)** %TNFα^+^ CD4^+^ T cells **C)** %IL10^+^ CD4^+^ T cells **D)** %FOXP3^+^CD4^+^ T cells **E)** %CD69^+^ CD4^+^ T cells (n≥3, *p*<.001***, *p*<.01**, One-way ANOVA).

### Effects of silencing csMHCII on the TME

To investigate PD-L1 expression in the TME we injected CMT167-NT-ctrl and CMT167-shCIITA cells into GFP+ mice, recovered the GFP+ and GFP-(negative) populations and analyzed PD-L1 expression by flow cytometry. The GFP-(negative) cancer cells had a decrease in PD-L1 expression *in vivo*: however there was no change in PD-L1 expression on the GFP+ cells, in the TME as a whole (**Figure 4A, 4B**). To further characterize the TME we used Spectral Flow Cytometry on the CYTEK Aurora to look at 22 markers. We harvested mice with CMT167-NT-ctrl tumors and CMT167-shCIITA tumors. MHCII expression on cells in the TME did not change (**Supplemental Figure S1J**); however we were interested to see how MHCII was expressed on specific antigen presenting cells. We analyzed the CYTEK data with unsupervised clustering analysis shown in a tSNE plot (**Figure 4C**). There were changes in the B cell clusters and some macrophage clusters but nothing reached statistical significance. To further analyze the data we used bi-axial gating to characterize the populations in the TME as previously defined [25, 28]. There was an increase in B cells in the CIITA knockdown tumors and these B cells had increased expression of PD-L1 and MHCII (**Figure 4D-4F**). In the CIITA knockdowns we detected no change in resident alveolar macrophages, but there was decrease in recruited macrophages. There was no change in macrophages exhaustion markers or PD-L1 expression, and a slight increase in MHCII expression (**Figure 4G-4J**). There was no change in dendritic cell or neutrophils recruitment: however the DC showed a slight increase in MHCII expression (**Figure 4K-4N**). These data suggest that in the setting of less antigen presentation by cancer cells, due to lack of csMHCII there is increased antigen presentation by B cells and other APC.

**Figure 4:**
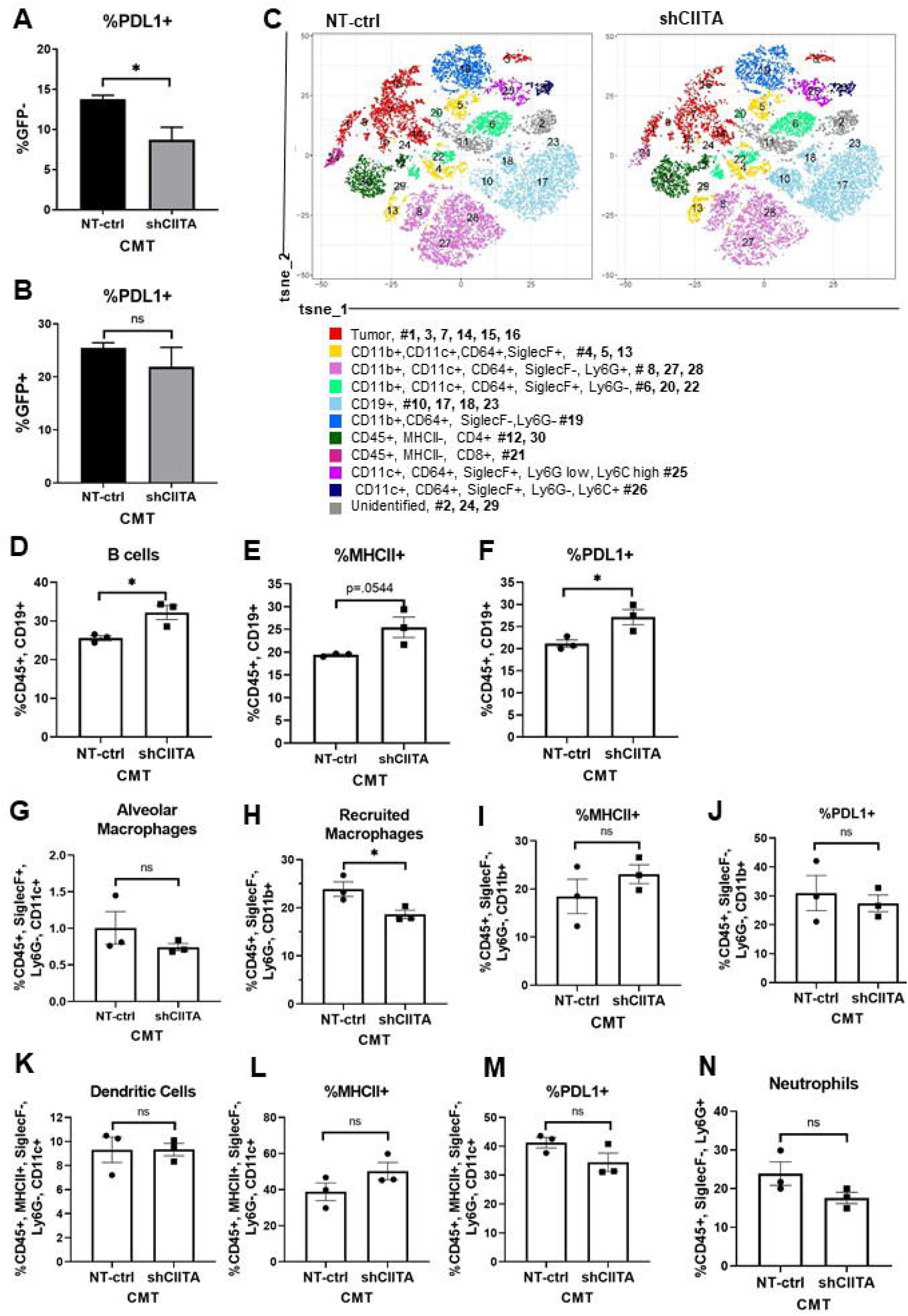
Effects of silencing csMHCII on the TME. Flow cytometric analysis of %PD-L1 of **A)** GFP-(neg) and **B)** GFP+ cells from CMT167-NT-ctrl and CMT167-shCIITA tumors injected into GFP+ mice and FACs sorted (n=3, p<.05*, Unpaired T-test). **C)** tSNE plot of unsupervised clustering analysis of Spectral Flow CYTEK of CMT167-NT-ctrl and CMT167-shCIITA. Bi-axial analysis of Spectral Flow CYTEK of **D)** B cells (CD45+, CD19+) **E)** MHCII+ B cells and **F)** PDL1+ B cells **G)** Alveolar Macrophages (CD45+, SiglecF+, Ly6G−, CD11c+) **H)** recruited macrophages (CD45+, SiglecF−, Ly6G−, CD11b+) **I)** MHCII+ Recruited macrophages **J)** PDL1+ recruited macrophages **K)** Dendritic Cells (CD45+, MHCII+, SiglecF−, Ly6G−, CD11c+) **L)** MHCII+ Dendritic Cells, **M)** PDL1+ Dendritic Cells, **N)** Neutrophils (n=3, p<.05*, Unpaired T test).

### Overexpression of CIITA in LLC cells induces MHCII and increases CD4^+^ and CD8^+^ tumor infiltration

Since loss of csMHCII in CMT167 resulted in decreased T cell infiltration into tumors, we sought to determine if induction of csMHCII in LLC cells, a cell line that does not express MHCII, would be sufficient to increase T cell infiltration. LLC cells were transfected with a lentiviral expression vector (LLC-CIITAOE) or an empty vector control (LLC-EV-ctrl). CIITA overexpression induced surface MHCII expression at baseline, and expression of MHCII associated genes (**Figure 5A 5B, and Supplemental Figure S3A, S3B**). Tumor-bearing lungs were stained for tumor infiltrating T cells. LLC-CIITAOE tumors had increased infiltration of CD3^+^ (EV-ctrl 36.35±12.3 vs. CIITAOE 89.93±8.766), CD4^+^ (EV-ctrl 19.48±9.595 vs. CIITAOE 37.41±7.881) and CD8^+^ T cells (EV-ctrl 17.26±3.014 vs. CIITAOE 59.75±22.46) (**Figure 5C**). Representative images for LLC-EV-ctrl and LLC-CIITAOE tumors are shown in (**Figure 5D**).

**Figure 5:**
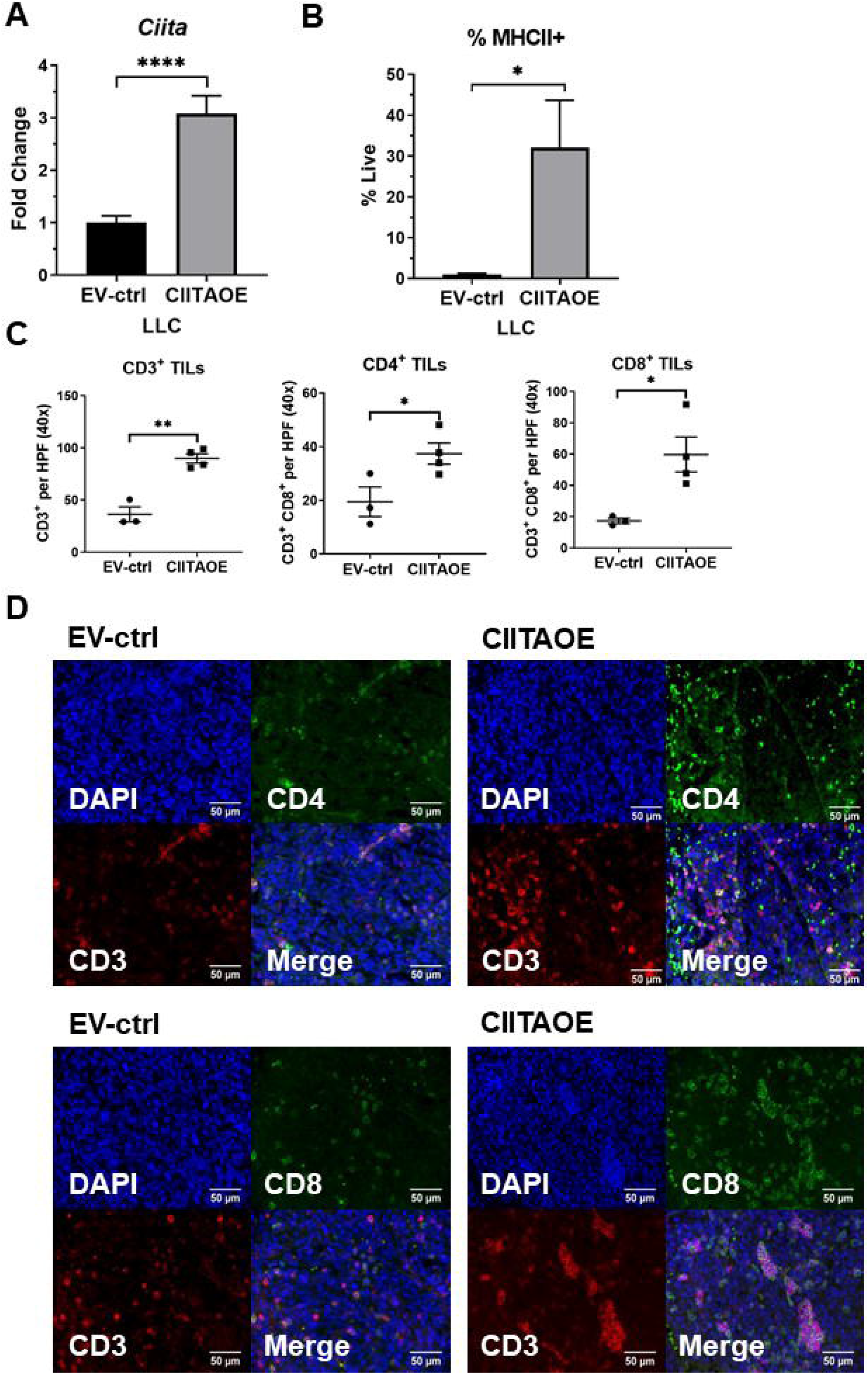
Overexpression of CIITA in LLC cells induces MHCII and increases CD4^+^ and CD8^+^ tumor infiltration. **A**) Fold change of *Ciita* mRNA expression for LLC-EV-ctrl and LLC-CIITAOE (n=3, p<.0001****, Unpaired T-test) **.B)** Flow Cytometry analysis of MHCII on LLC-EV-ctrl and LLC-CIITAOE cells (n=3, p<.05*, Unpaired T-test). **C)** Average Count of immunofluorescence of CD3^+^, CD4^+^, and CD8^+^ positive T cells per high power field (40x) in LLC-EV-ctrl tumors and LLC-CIITAOE tumors (n=3, n=4, *p*<.05*, *p*<.01**, Unpaired T-test). **D)** Representative pictures of LLC-EV-ctrl and LLC-CIITAOE tumors IF, DAPI (blue), CD3 (red), and CD4 or CD8 (green).

### Effects of csMHCII expression on tumor progression and response to anti-PD-1

To determine the effect of silencing csMHCII on tumor progression and response to immunotherapy, equal numbers of CMT167-NT-ctrl or CMT167-shCIITA cells were injected into the lungs of wild-type mice, and primary tumor size was measured at 4 weeks. Compared to CMT167-NT-ctrl tumors, the CMT167-shCIITA tumors had an increase in tumor size (NT-ctrl 15.07mm^3^±1.674 vs. shCIITA 21.37mm^3^±1.821) (**Figure 6A**). Separate groups of mice were treated starting 7 days after tumor implantation with anti-PD-1 antibody or IgG isotype control as previously described [24]. Anti-PD-1 therapy resulted in almost complete ablation of CMT167-NT-ctrl tumors, similar to what we have previously shown with parental CMT167 tumors [24]. In contrast, CMT167-shCIITA tumors had a diminished response to anti-PD-1 therapy, with only a 50% inhibition of tumor growth in response to anti-PD-1 (**Figure 6B**). These data demonstrate that csMHCII is necessary for optimal response to anti-PD-1 therapy. Thus, loss of csMHCII expression increases tumor progression, and at least partially reverses responsiveness to anti-PD-1.

**Figure 6:**
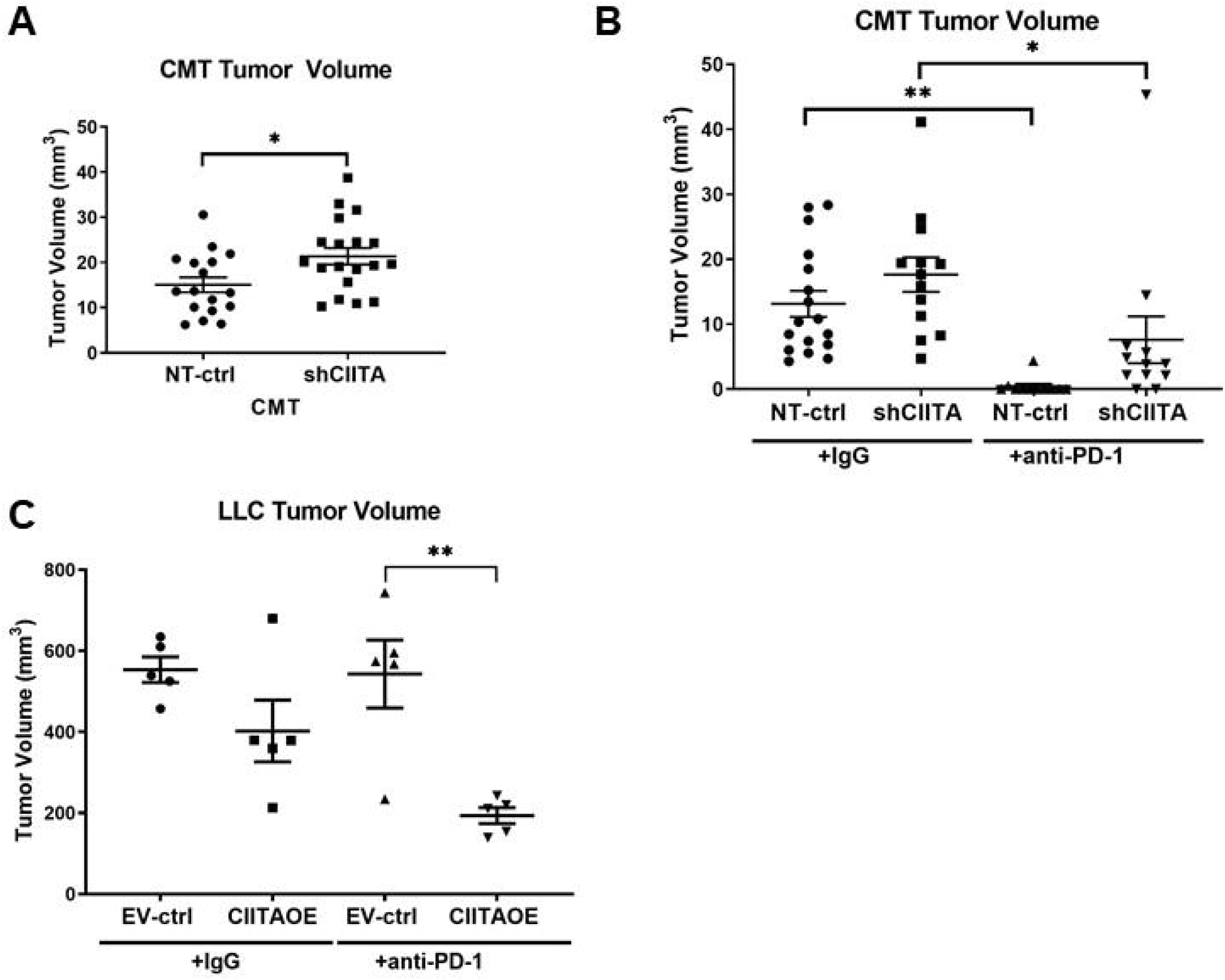
MHCII mediates the tumor progression and the response to checkpoints. **A)** Primary tumor volume of CMT167-NT-ctrl or CMT167-shCIITA tumors at 4 weeks (n≥17, *p*<.05*, Unpaired T-test). **B)** Primary tumor volume of CMT167-NT-ctrl or CMT167-shCIITA treated with IgG antibody or PD-1 antibody at 4.5 weeks (n≥11, *p*<.01**, Two-way ANOVA). **C)** Primary tumor volume of LLC-EV-ctrl or LLC-CIITAOE treated with IgG antibody at 3 weeks (n=5, *p*<.01**, Two-way ANOVA).

To determine if induction of csMHCII in LLC cells would be sufficient to sensitize the tumors to anti-PD1 *in vivo*, LLC EV-ctrl or CIITAOE cells were injected into C57BL/6 WT mice. Tumors were allowed to develop for 7 days and then mice were treated with either an IgG control or anti-PD-1 antibody for 2 weeks. In mice injected with control antibody, LLC-EV-ctrl and LLC-CIITAOE tumors had comparable growth, indicating that enforced csMHCII expression was not sufficient to alter tumorigenesis under baseline conditions *in vivo*. However, while anti-PD-1 antibody had no impact on tumorigenesis in mice injected with LLC-EV-ctrl cells, we observed ~70% decrease in primary tumor volume in mice implanted with LLC-CIITAOE tumor cells (**Figure 6C**). In total, these data demonstrate that csMHCII can regulate responses to anti-PD-1 therapy in mouse models of NSCLC.

### Adenocarcinoma Lung Cancer expresses HLA-DR (MHCII) and is correlated with enhanced CD3^+^, CD4^+^ and CD8^+^ TILs in human lung tumors

To extend our findings on csMHCII to human lung cancer, we quantified MHCI and MHCII expression in a panel of human NSCLC cell lines using RNA-seq data from the Cancer Cell Line Encyclopedia (CCLE) for 15 KRAS mutant lung cancer cell lines [27]. MHCI mRNA expression (HLA-A, HLA-B, HLA-C), and MHCI accessory genes were high across almost all lung adenocarcinoma cell lines (**Supplemental Figure S3C**). However, MHCII mRNA expression (HLA-DR, HLA-DB, and HLA-DQ) and MHCII accessory genes were only expressed in a subset of cell lines (**Supplemental Figure S3D**). We validated HLA-DR, HLA-DQ, and CD74 protein expression in a subset of these human lung cancer cell lines using flow cytometry (**Supplemental Figure S3E**). The cell lines could be divided into 3 subsets: 1)high basal levels; 2)low basal levels which were not altered by IFNγ; and 3)low levels with the ability to induce MHCII in response to IFNγ. H1155 had low basal levels and did not induce HLA-DR, A549 has low basal levels and induced HLA-DR, and H1373 had high basal levels with the ability to induce HLA-DR (**Supplemental Figure S3F**).

In addition to characterizing MHCII expression in human lung cancer cell lines, we quantified csMHCII in primary human lung adenocarcinomas. To do this, we applied multispectral imaging using the Vectra 3.0 system to quantify csMHCII in a panel of 90, resected human lung adenocarcinomas patients using the Vectra 3.0 instrument. To define csMHCII expression, we stained tissue sections for cytokeratin (CK), marking the cancer cells, and HLA-DR, a human MHCII molecule, and analyzed the incidence of double positive staining within the tumor. Using an arbitrary cutoff we defined csMHCII+ positive tumors as those exhibiting ≥ 10% double positive cells. Cancer-specific positive HLA-DR (≥10%) was observed in 51 of the 90 tumors investigated (57%), consistent with previous findings [31]. Patients with HLA-DR positive tumors (≥10%) had a 5 year overall survival benefit (**Figure 7A**). Tumor cell expression of HLA-DR+ events were positively correlated with, CD4^+^ events in the tumor (**Figure 7B**). In addition HLA-DR positive tumors (≥10%) had an increase in CD4^+^ T cell infiltration and a slight increase in CD8^+^ T cell infiltration although it did not reach statistical significance, recapitulating our results in the murnine model (**Figure7C and 7D**). Representative images show DAPI (blue), CK (magenta), HLA-DR (yellow), CD3 (green), and CD8 (red) of a HLA-DR negative tumor and an HLA-DR positive tumor (**Figure 7E and 7F respectively**). These data demonstrate that csMHCII is a feature of many human lung adenocarcinomas, and that csMHCII is positively associated with T cell infiltration, strongly suggesting the potential value of csMHCII as a potential biomarker that may facilitate better patient stratification for response to immunotherapy.

**Figure 7:**
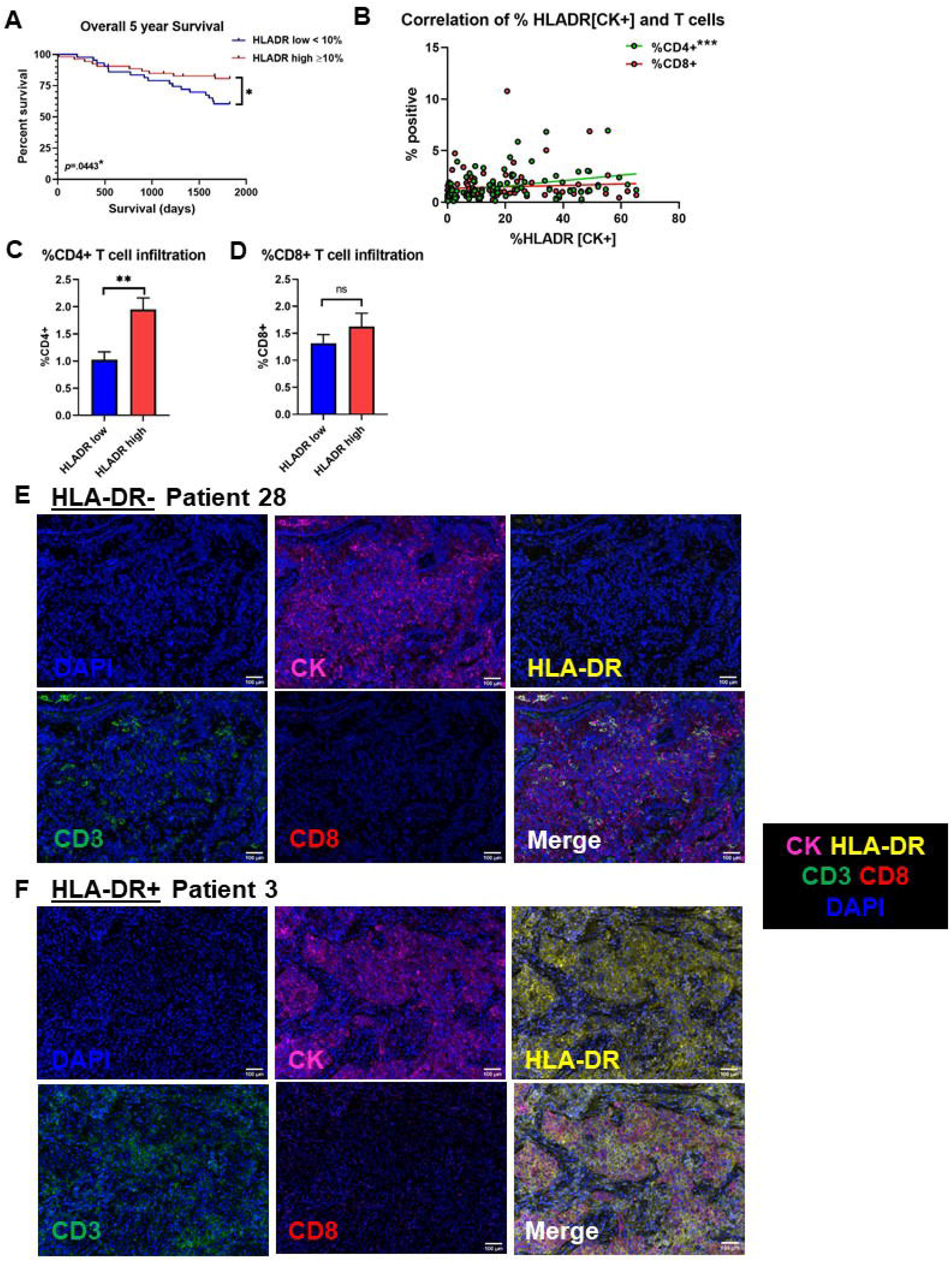
Adenocarcinoma Lung Cancer expresses HLA-DR (MHCII) and is correlated with enhanced CD3^+^, CD4^+^ and CD8^+^ TILs. **A)** Overall 5 year Survival Curve stratified by HLADR low <10% and HLADR high ≥10% patients (n=90, *p*<.05*, Log-rank (Mantel-Cox) test) **B)** The correlation of the percent of the number of counts of HLADR+ on cancer cells and the percent of the number of counts of CD8 (r^2^=.0069, *p*=.439), CD4 (CD3+, CD8-) (r^2^=.1227, *p*<.001***) in the tumor. **C)** %CD4+ and **D)** CD8+ T cell infiltration in the tumor in patients stratified by HLADR low <10% and HLADR high ≥10% (n=90, p<.001**, Unpaired T-test). **E)** Representative images of the composite image after spectral unmixing. DAPI nuclear marker (pseudocolored blue), Cytokeratin (CK) (membrane, Cy3, pseudocolored magenta), HLA-DR (membrane, Cy5, pseudocolored yellow), CD3 (membrane, FITC, pseudocolored green), CD8 (membrane, Texas Red, pseudocolored red) of **D)** HLA-DR-tumor of patient 28 and **E)** HLA-DR+ tumor of patient 3.

## Discussion

The ability of the immune system to recognize antigens is critical for tumor elimination by T cells. Classically, this is mediated through presentation of antigens via MHCI expressed on cancer cells and MHCII expressed on professional APCs. However, non-APCs, such as endothelial cells and epithelial cells, including lung epithelial cells, can also express MHCII [32–35]. These cells induce MHCII expression in response to IFNγ, with the capability to directly present antigen to CD4^+^ T cells [36, 37]. While the potential impact of MHCII expression on anti-tumor immunity has been appreciated for years, recent studies have provided new evidence for the ability of MHCII to influence the mutational landscape of diverse tumor types [38]. Furthermore, MHCII expression on cancer cells has been observed in a variety of cancers and in general correlates with improved outcomes. In colon cancer, high csMHCII expression is associated with increased survival [18]. In melanoma, csMHCII expression is associated with a better response to checkpoint therapy blockade with an increase in progression-free and overall survival in anti-PD-1/PD-L1 treated patients stratified by csMHCII positivity [20]. In breast cancer, csMHCII expression is downregulated as a consequence of decreased CIITA in highly metastatic cells, and in TNBC tumor-specific MHCII is associated with increased progression free survival [22, 39].

Our interest on csMHCII began with the finding that in our mouse model csMHCII was correlated with response to immunotherapy, with anti-PD-1-sensitive CMT167 tumors characterized by csMHCII expression and anti-PD-1-resistant LLC tumors failing to express csMHCII. A major goal of these studies was to identify whether csMHCII had a functional consequence on shaping the TME and/or the response of tumors to immunotherapy. Silencing CIITA, and in turn csMHCII, in CMT167 cells resulted in slightly larger tumors under baseline conditions. CIITA knockdown was further associated with reduced CD4^+^ helper T cell infiltration and Th1 cytokine production, as well as IFNγ expressing CD8^+^ T cells. This was not a consequence of alterations in MHCI on the cancer cells. Using spectral flow antibody staining we have demonstrated that silencing CIITA causes more global changes in the TME. We detected decreases in recruited macrophage populations and increases in B cells. Importantly, silencing CIITA in cancer cells resulted in decreased production in the TME of CXCL9, a chemokine critical for T cell recruitment. Consistent with decreasing T cell infiltration into the tumors, csMHCII had a pronounced role in regulating the response to anti-PD-1 targeted immunotherapy in CMT167 tumors; inhibiting CIITA and csMHCII expression converted an immunotherapy-sensitive tumor to an immunotherapy-resistant phenotype. Conversely, expression of CIITA and csMHCII in LLC cells increased T cell infiltration into these tumors, and converted them from immunotherapy-resistant to immunotherapy-sensitive.

A critical consequence of csMHCII is the ability of cancer cells to directly present tumor neoantigens to CD4^+^ T cells, providing an APC-independent mechanism for CD4^+^ T cell activation in the TME. The capacity of cancer cells to directly activate CD4^+^ T cells was confirmed by *in vitro* co-culture studies of cancer cells and T cells, where cancer cells lacking csMHCII expression showed a significant decrease in effector cytokine production of Th1 CD4^+^ T cells compared to cancer cells expressing MHCII. The identity of tumor-specific MHCII neoantigens remains unclear at this time, but one likely source for these peptides is likely to arise from autophagy, a cellular process associated with the generation of MHCII epitopes [40, 41]. It is also possible that neoantigens presented by the cancer cells may be distinct from those presented by MHCII on professional APCs in the TME. The unique contribution of MHCII is very likely contributed by the distinct peptide binding characteristics of MHCII compared to MHCI [42], with MHCII peptide presentation linked to tumor control in different contexts [43]. A recent study has examined specific CD4+ T cells neoantigens in human lung tumors, and while most CD4^+^ T cells recognized passenger mutations, a subset recognized the oncogenic driver [44]. The status of csMHCII was not examined in that study. Further investigation of the diversity of T cell receptors elicited in the presence or absence of csMHCII will be particularly informative in defining the impact of csMHCII on antitumor immunity.

These data implicate csMHCII as a modifier of the TME, promoting T cell infiltration. Our results are consistent with a model in which direct interactions between cancer cells expressing MHCII and resident CD4^+^ T cells results in CD4^+^ activation leading to local IFNγ expression in the TME. This in turn leads to IFNγ-dependent changes including enhanced CXCL9 production and PD-L1 expression, which in turn causes further T cell recruitment. This positive feedback mechanism culminates in a T cell-inflamed, immunotherapy-sensitive tumor phenotype [24]. It has been proposed that in some cases cancer immunotherapies achieve limited results due to a lack of successful activation of anti-tumor CD4^+^ T cells [45, 46]. Specifically, Th1 CD4^+^ T cells can play a crucial role in the tumor microenvironment, facilitating optimal CD8^+^ T cell activation and effector function. Th1 cytokines, including IFNγ and TNFα, can further induce cancer cell senescence [47]. While other factors, such as mutational burden and presence of neoantigens contribute to T cell infiltration and response to immunotherapy [48], in these experiments we used isogenic cells in which only the expression of CIITA has been altered. Thus, our data support a model for csMHCII as biomarker of sensitivity to checkpoint inhibitors independent of other factors, including MHCII expression by other cells in the TME. Although csMHCII expression has been reported on NSCLC and associated with lymphocytic infiltration [31, 49], this was not previously correlated with response to immunotherapy. Our data show that csMHCII is expressed in a subset of unselected human lung adenocarcinomas and is correlated with an increase in T cell tumor infiltration and a 5 year overall survival benefit. However oncogenic drivers were not defined for many of these patients. Additional studies on a larger cohort in which oncogenic drivers have been defined, and where response to immunotherapy can be correlated are necessary to better define effects of csMHCII on survival and response to immunotherapy. Our studies provide direct evidence for the functional importance of MHCII expression on cancer cells, and illustrate that MHCII expression on professional APCs, including dendritic cells, is not sufficient to confer susceptibility to immunotherapy.

