## Supplemental for "Cancer Cell Intrinsic Expression of MHCII Regulates the Immune Microenvironment and Response to Anti-PD-1 Therapy in Lung Adenocarcinoma"

### Slide 1
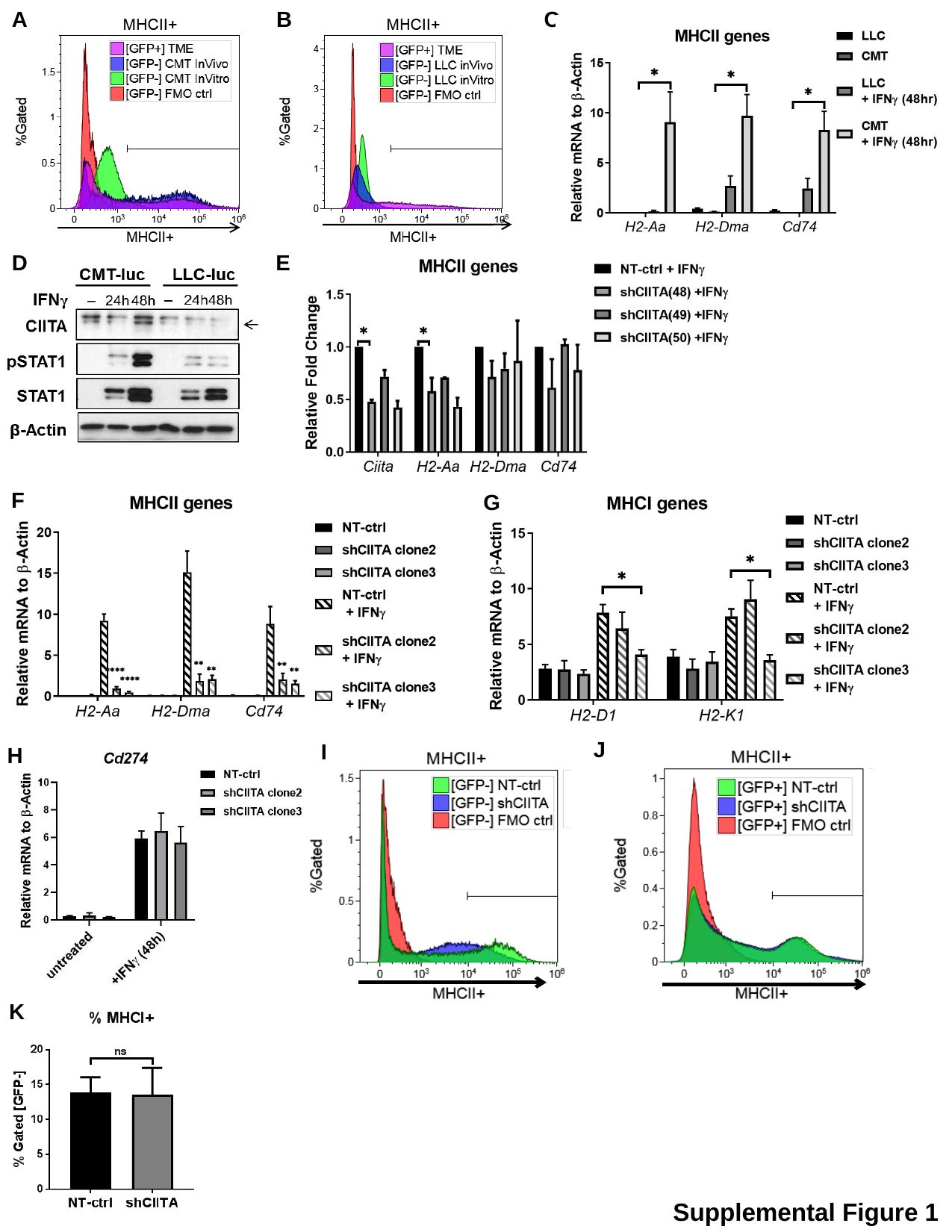

A
B
C
D
E
F
G
J
I
H
K
Supplemental Figure 1

### Slide 2
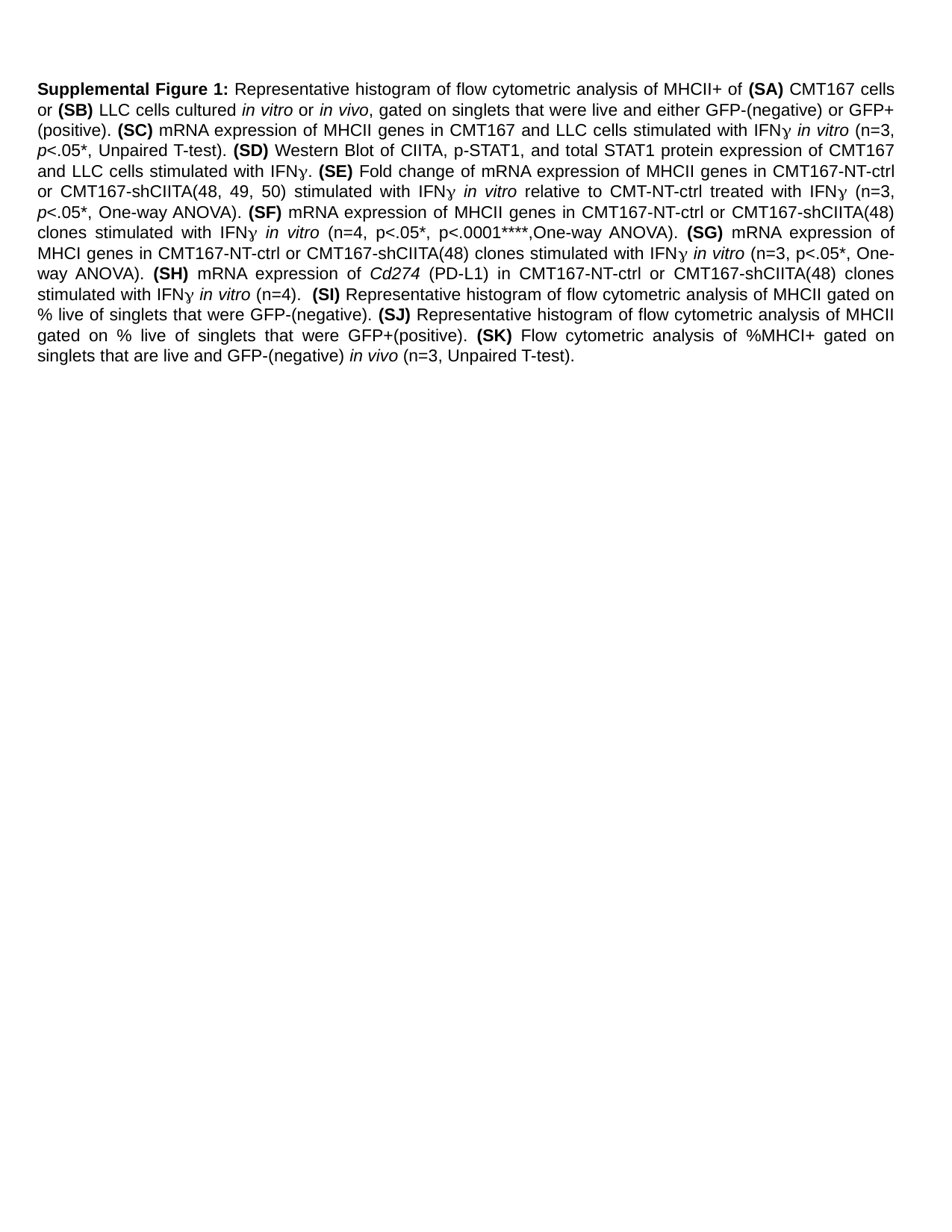

Supplemental Figure 1: Representative histogram of flow cytometric analysis of MHCII+ of (SA) CMT167 cells or (SB) LLC cells cultured in vitro or in vivo, gated on singlets that were live and either GFP-(negative) or GFP+(positive). (SC) mRNA expression of MHCII genes in CMT167 and LLC cells stimulated with IFNγ in vitro (n=3, p<.05*, Unpaired T-test). (SD) Western Blot of CIITA, p-STAT1, and total STAT1 protein expression of CMT167 and LLC cells stimulated with IFNγ. (SE) Fold change of mRNA expression of MHCII genes in CMT167-NT-ctrl or CMT167-shCIITA(48, 49, 50) stimulated with IFNγ in vitro relative to CMT-NT-ctrl treated with IFNγ (n=3, p<.05*, One-way ANOVA). (SF) mRNA expression of MHCII genes in CMT167-NT-ctrl or CMT167-shCIITA(48) clones stimulated with IFNγ in vitro (n=4, p<.05*, p<.0001****,One-way ANOVA). (SG) mRNA expression of MHCI genes in CMT167-NT-ctrl or CMT167-shCIITA(48) clones stimulated with IFNγ in vitro (n=3, p<.05*, One-way ANOVA). (SH) mRNA expression of Cd274 (PD-L1) in CMT167-NT-ctrl or CMT167-shCIITA(48) clones stimulated with IFNγ in vitro (n=4). (SI) Representative histogram of flow cytometric analysis of MHCII gated on % live of singlets that were GFP-(negative). (SJ) Representative histogram of flow cytometric analysis of MHCII gated on % live of singlets that were GFP+(positive). (SK) Flow cytometric analysis of %MHCI+ gated on singlets that are live and GFP-(negative) in vivo (n=3, Unpaired T-test).

### Slide 3
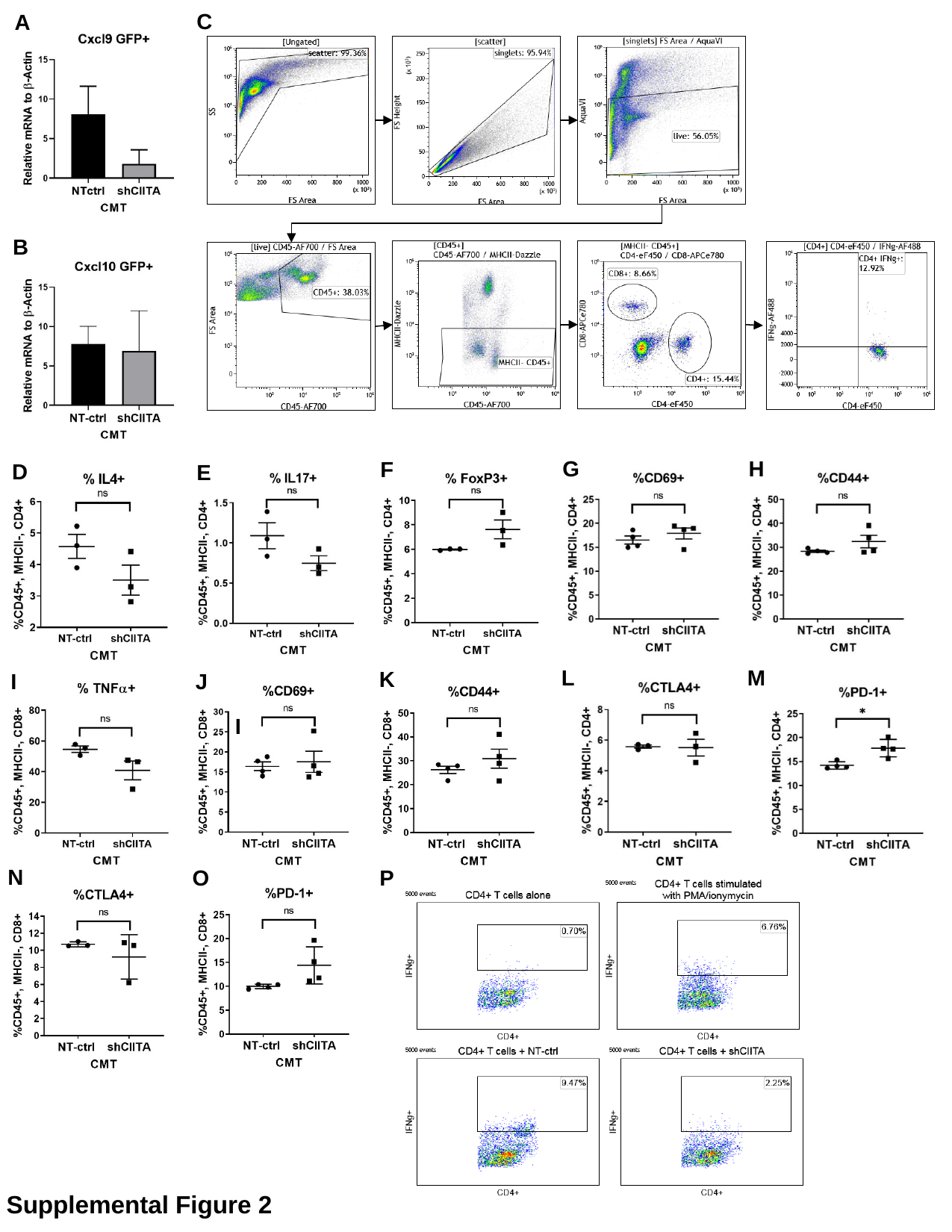

C
A
B
G
H
F
D
E
Supplemental Figure 2: qRT-pcr on GFP+ cells FACs sorted from CMT167-NT-ctrl and CMT167-shCIITA for SA) cxcl9, and SB) cxcl10. SC) Gating strategy for T cell flow cytometry, events were ganed on singlets that were live that were CD45+ and MHCII- and either CD4+ or CD8+. Flow cytometry on tumor bearing lungs of the frequency of T cells compared between mice injected with CMT167-NT-ctrl and CMT167-shCIITA tumors (n=4, Unpaired T-test) of CD4 T cells that were D) IL4+, E) IL17+, F) FoxP3+, G)CD69+, H) CD44+ or CD8 T cells I)TNFa+, J) CD69+, and K) CD44+ . Flow Cytometry analysis of exhaustion markers on CD4 T cells L) PD1 and M) CTLA4 and CD8 T cells N) CTLA4, and O) PD1. Flow Cytometry Plot for CD4^+^ T cells gated on singlets that are live and CD45+. (SP) IFNγ^+^ for CD4^+^ T cells cultured alone, CD4^+^ T cells stimulated with PMA/ionomycin, CD4^+^ T cells co-cultured with CMT-NT-ctrl cells, and CD4^+^ T cells co-cultured with CMT-shCIITA.
L
M
K
I
J
I
N
O
P
J
K
Supplemental Figure 2

### Slide 4
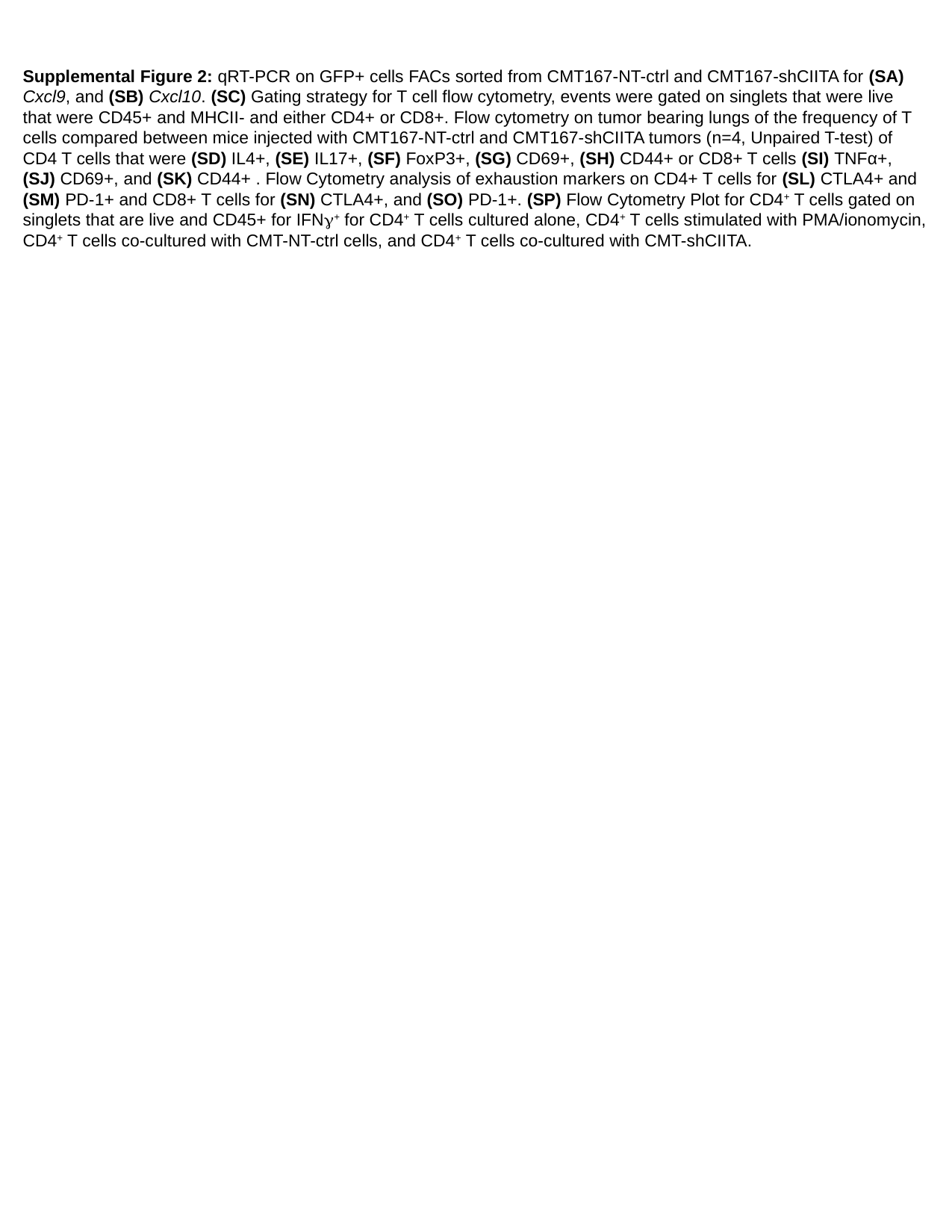

Supplemental Figure 2: qRT-PCR on GFP+ cells FACs sorted from CMT167-NT-ctrl and CMT167-shCIITA for (SA) Cxcl9, and (SB) Cxcl10. (SC) Gating strategy for T cell flow cytometry, events were gated on singlets that were live that were CD45+ and MHCII- and either CD4+ or CD8+. Flow cytometry on tumor bearing lungs of the frequency of T cells compared between mice injected with CMT167-NT-ctrl and CMT167-shCIITA tumors (n=4, Unpaired T-test) of CD4 T cells that were (SD) IL4+, (SE) IL17+, (SF) FoxP3+, (SG) CD69+, (SH) CD44+ or CD8+ T cells (SI) TNFα+, (SJ) CD69+, and (SK) CD44+ . Flow Cytometry analysis of exhaustion markers on CD4+ T cells for (SL) CTLA4+ and (SM) PD-1+ and CD8+ T cells for (SN) CTLA4+, and (SO) PD-1+. (SP) Flow Cytometry Plot for CD4^+^ T cells gated on singlets that are live and CD45+ for IFNγ^+^ for CD4^+^ T cells cultured alone, CD4^+^ T cells stimulated with PMA/ionomycin, CD4^+^ T cells co-cultured with CMT-NT-ctrl cells, and CD4^+^ T cells co-cultured with CMT-shCIITA.

### Slide 5
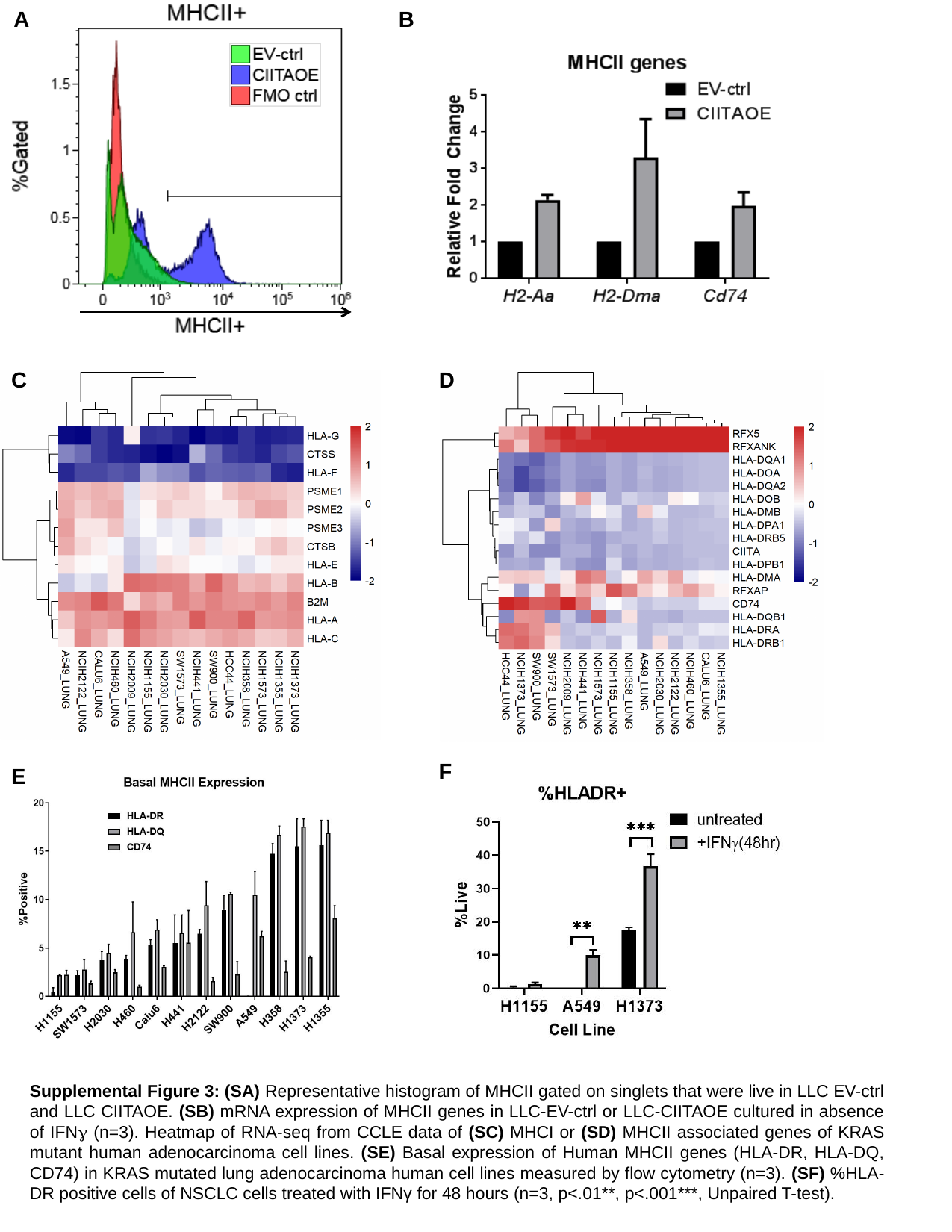

A
B
C
D
F
E
Supplemental Figure 3: (SA) Representative histogram of MHCII gated on singlets that were live in LLC EV-ctrl and LLC CIITAOE. (SB) mRNA expression of MHCII genes in LLC-EV-ctrl or LLC-CIITAOE cultured in absence of IFNγ (n=3). Heatmap of RNA-seq from CCLE data of (SC) MHCI or (SD) MHCII associated genes of KRAS mutant human adenocarcinoma cell lines. (SE) Basal expression of Human MHCII genes (HLA-DR, HLA-DQ, CD74) in KRAS mutated lung adenocarcinoma human cell lines measured by flow cytometry (n=3). (SF) %HLA-DR positive cells of NSCLC cells treated with IFNγ for 48 hours (n=3, p<.01**, p<.001***, Unpaired T-test).

### Slide 6
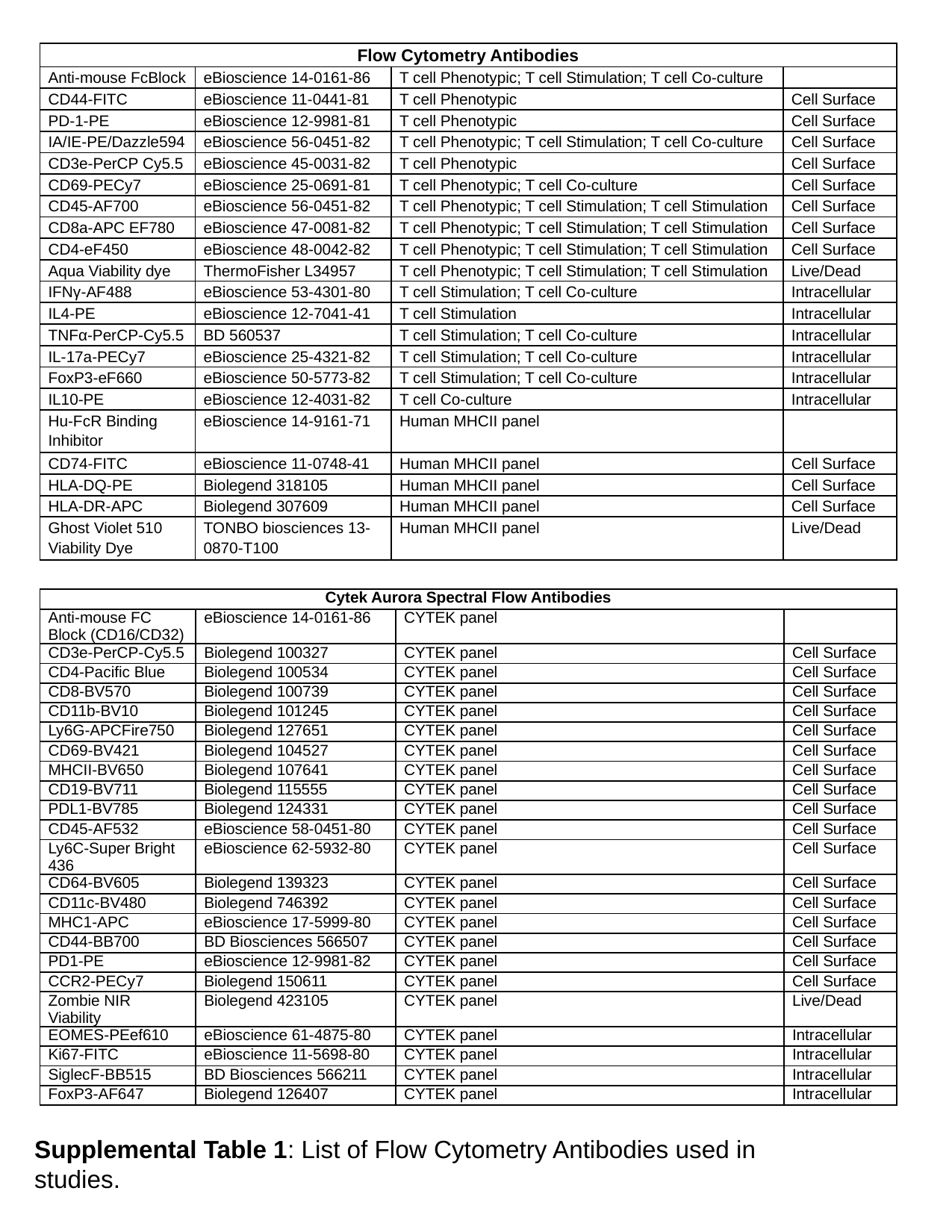

| Flow Cytometry Antibodies | | | |
| --- | --- | --- | --- |
| Anti-mouse FcBlock | eBioscience 14-0161-86 | T cell Phenotypic; T cell Stimulation; T cell Co-culture | |
| CD44-FITC | eBioscience 11-0441-81 | T cell Phenotypic | Cell Surface |
| PD-1-PE | eBioscience 12-9981-81 | T cell Phenotypic | Cell Surface |
| IA/IE-PE/Dazzle594 | eBioscience 56-0451-82 | T cell Phenotypic; T cell Stimulation; T cell Co-culture | Cell Surface |
| CD3e-PerCP Cy5.5 | eBioscience 45-0031-82 | T cell Phenotypic | Cell Surface |
| CD69-PECy7 | eBioscience 25-0691-81 | T cell Phenotypic; T cell Co-culture | Cell Surface |
| CD45-AF700 | eBioscience 56-0451-82 | T cell Phenotypic; T cell Stimulation; T cell Stimulation | Cell Surface |
| CD8a-APC EF780 | eBioscience 47-0081-82 | T cell Phenotypic; T cell Stimulation; T cell Stimulation | Cell Surface |
| CD4-eF450 | eBioscience 48-0042-82 | T cell Phenotypic; T cell Stimulation; T cell Stimulation | Cell Surface |
| Aqua Viability dye | ThermoFisher L34957 | T cell Phenotypic; T cell Stimulation; T cell Stimulation | Live/Dead |
| IFNγ-AF488 | eBioscience 53-4301-80 | T cell Stimulation; T cell Co-culture | Intracellular |
| IL4-PE | eBioscience 12-7041-41 | T cell Stimulation | Intracellular |
| TNFα-PerCP-Cy5.5 | BD 560537 | T cell Stimulation; T cell Co-culture | Intracellular |
| IL-17a-PECy7 | eBioscience 25-4321-82 | T cell Stimulation; T cell Co-culture | Intracellular |
| FoxP3-eF660 | eBioscience 50-5773-82 | T cell Stimulation; T cell Co-culture | Intracellular |
| IL10-PE | eBioscience 12-4031-82 | T cell Co-culture | Intracellular |
| Hu-FcR Binding Inhibitor | eBioscience 14-9161-71 | Human MHCII panel | |
| CD74-FITC | eBioscience 11-0748-41 | Human MHCII panel | Cell Surface |
| HLA-DQ-PE | Biolegend 318105 | Human MHCII panel | Cell Surface |
| HLA-DR-APC | Biolegend 307609 | Human MHCII panel | Cell Surface |
| Ghost Violet 510 Viability Dye | TONBO biosciences 13-0870-T100 | Human MHCII panel | Live/Dead |
| Cytek Aurora Spectral Flow Antibodies | | | |
| --- | --- | --- | --- |
| Anti-mouse FC Block (CD16/CD32) | eBioscience 14-0161-86 | CYTEK panel | |
| CD3e-PerCP-Cy5.5 | Biolegend 100327 | CYTEK panel | Cell Surface |
| CD4-Pacific Blue | Biolegend 100534 | CYTEK panel | Cell Surface |
| CD8-BV570 | Biolegend 100739 | CYTEK panel | Cell Surface |
| CD11b-BV10 | Biolegend 101245 | CYTEK panel | Cell Surface |
| Ly6G-APCFire750 | Biolegend 127651 | CYTEK panel | Cell Surface |
| CD69-BV421 | Biolegend 104527 | CYTEK panel | Cell Surface |
| MHCII-BV650 | Biolegend 107641 | CYTEK panel | Cell Surface |
| CD19-BV711 | Biolegend 115555 | CYTEK panel | Cell Surface |
| PDL1-BV785 | Biolegend 124331 | CYTEK panel | Cell Surface |
| CD45-AF532 | eBioscience 58-0451-80 | CYTEK panel | Cell Surface |
| Ly6C-Super Bright 436 | eBioscience 62-5932-80 | CYTEK panel | Cell Surface |
| CD64-BV605 | Biolegend 139323 | CYTEK panel | Cell Surface |
| CD11c-BV480 | Biolegend 746392 | CYTEK panel | Cell Surface |
| MHC1-APC | eBioscience 17-5999-80 | CYTEK panel | Cell Surface |
| CD44-BB700 | BD Biosciences 566507 | CYTEK panel | Cell Surface |
| PD1-PE | eBioscience 12-9981-82 | CYTEK panel | Cell Surface |
| CCR2-PECy7 | Biolegend 150611 | CYTEK panel | Cell Surface |
| Zombie NIR Viability | Biolegend 423105 | CYTEK panel | Live/Dead |
| EOMES-PEef610 | eBioscience 61-4875-80 | CYTEK panel | Intracellular |
| Ki67-FITC | eBioscience 11-5698-80 | CYTEK panel | Intracellular |
| SiglecF-BB515 | BD Biosciences 566211 | CYTEK panel | Intracellular |
| FoxP3-AF647 | Biolegend 126407 | CYTEK panel | Intracellular |
Supplemental Table 1: List of Flow Cytometry Antibodies used in studies.
